# Targeted intervention of senescence induced by a 5′ half fragment of tRNA^Seca(NCA)^ with antisense oligonucleotide extends healthspan and lifespan in mice

**DOI:** 10.1101/2025.03.24.644848

**Authors:** Kai-Yue Cao, Long-Bo Bai, Da Zhang, Yu-Hang Zhang, Rui-Ze Gong, Tong-Meng Yan, Yu Pan, Ya-Ting Cheng, Zhi-Hong Jiang

## Abstract

In light of increasing attention being paid to aging research globally, the accumulation of aging hallmarks and their corresponding targeting therapeutics have been substantially revealed. However, uncovering the genuine drivers within epigenetic alterations that lead to aging remains a formidable challenge. In this study, we identified tRNA^Sec(NCA)^ as the most severely damaged tRNA species in the kidneys of naturally aged mice. This damage not only dysregulated selenoproteins with anti-aging effects, but also generates a 5′-tRNA fragment cleaved at the 34^th^ position, which accumulates in an age-dependent manner. Mechanistically, the 5′-tRNA^Sec(NCA)^ half interacts and activates Toll-like receptor 7, thereby triggering innate immune responses and promoting cellular senescence in both mice and human cells. Moreover, in a naturally aged mice model, administration of an antisense oligonucleotide (ASO) targeting the 5′-tRNA^Sec(NCA)^ half remarkably ameliorates aging markers, enhances telomere length, and extends healthspan and lifespan. In addition, ASO-5′-tRNA^Sec(NCA)^ half plays another role in directly targeting and downregulating BCAT1 *via* the RNAi pathway to intervene in the senescence process. Our findings underscore tRNA damage as a novel aging hallmark, and targeting the damage-induced products presents a novel strategy for aging intervention, thus expanding our knowledge of the aging process.

## Introduction

Aging represents a significant global challenge, impacting nearly every aspect of human health and society. By the late 2070s, the number of people aged 65 years or older is projected to reach 2.2 billion, up from 962 million people aged 60 years or older in 2017, reflecting a dramatic demographic shift^1^. This increase in the aging population is accompanied by a surge in the prevalence of age-related diseases such as cardiovascular disorders, neurodegenerative diseases, and metabolic syndromes, which collectively contribute to over 70% of global deaths^2^. These conditions not only reduce the quality of life for millions of individuals but also impose substantial economic burdens on healthcare systems worldwide. While extending lifespan has been a focal point of scientific inquiry, the concept of “healthspan”-the period of life lived in good health, free from the chronic diseases and disabilities of aging-has increasingly come into focus^3^. Advances in biotechnology and molecular biology have begun to unravel the complex mechanisms underlying aging, offering promising avenues for interventions aimed at improving healthspan rather than merely prolonging life^3–5^. The shift toward enhancing healthspan emphasizes the importance of maintaining physical, cognitive, and emotional well-being in the elderly, thereby reducing the societal and economic impacts of aging^4^.

Therapeutic agents such as metformin^6^, rapamycin^7^, NAD^+^ precursors^8^, spermidine^9^, taurine^10^, senolytics^11^, GLP-1 agonists^12^ have been shown to slow aging process and extend lifespan in worms, flies, and mice, and some of these substances have entered clinical trials for validation. However, the development of anti-aging therapeutics still faces significant scientific and translational challenges. For instance, the innate complexity of aging biology, with its interlinked hallmarks from genomic instability to transcriptomic dysregulation, complicates targeted interventions since one pathway modulation may inadvertently disrupt compensatory mechanisms. Current small-molecule candidates struggle to balance efficacy with safety, exhibiting off-target effects, dose-dependent toxicity (e.g., immunosuppression in mTOR inhibitors), and limited tissue specificity in senolytic strategies^13^. Therefore, it is of crucial importance to explore the genuine incentives that drive aging, such as genomic instability, mitochondrial dysfunction, and chronic inflammation. Development of targeted anti-aging strategies based on these identified mechanisms holds greater promise in fundamentally intervening in the aging process and achieving more effective results compared to merely focusing on aging biomarkers^14^.

Epigenetic alterations, including DNA methylation^15^, histone modification^16^, chromatin remodeling^17^, and particularly non-coding RNAs (ncRNAs) regulation^18^, have been emerged as critical aging hallmarks. These changes influence gene expression without altering DNA sequence, significantly impacting cellular behavior during aging. In particular, as the most abundant small RNA species, tRNA has been addressed as crucial regulator of epigenetic alterations in aging^19^. The transcriptional activity of tRNA genes varies by cell types and changes during aging^20^. Meanwhile, the decline in protein synthesis rate during aging is closely related to the functional changes of tRNA^21^. It has been revealed that tRNA genes exhibit age-related DNA hypermethylation, which affects their expression and function^22^. The levels of tRNA-derived small RNAs in serum change with age and they are involved in regulating cellular stress responses, inflammatory responses, and other processes^23,24^. Unfortunately, the mechanisms by which tRNA expressions influence aging remain largely unexplored.

Previous multi-omics analyses have provided preliminary insights into the biological functions of transfer RNA-derived small RNAs (tsRNAs) in organ aging. Transcriptome RNA-seq revealed dynamic tsRNA expression patterns during early injury phases of skeletal muscle, demonstrating that upregulated 5′tiRNA-Gly post-injury promotes inflammatory responses and macrophage M1 polarization. This tsRNA species was found to regulate TGF-β signaling through targeting Tgfbr1, thereby enhancing the differentiation of muscle satellite cells and myocytes^24^. Meanwhile, RNA-seq identified elevated Glu-tsRNA-CTC levels in aged brains, which impairs mitochondrial translation and cristae organization by competitively binding to LaRs2 and inhibiting mt-tRNA^Leu^ aminoacylation, ultimately leading to reduced glutamate biosynthesis (GLS-dependent) and consequent synaptic structural defects and memory decline^25^. Intracerebroventricular administration of antisense oligonucleotides (ASOs) targeting Glu-tsRNA-CTC significantly improved mitochondrial translation and cristae defects, increased glutamate levels, rescued synaptic organizational deficits, and markedly enhanced memory function in aged mice across various behavioral paradigms. However, while the above study has revealed the therapeutic efficacy of targeting tsRNAs in mitigating Alzheimer’s disease-associated pathologies, whether targeted intervention against tsRNAs that promotes aging can effectively extend both healthspan and overall lifespan remains unexplored.

In the present study, multi-omics LC-MS technique was utilized to conduct a comprehensive profiling of tRNAs and proteins extracted from the kidneys of adult and aged mice, demonstrating that tRNA^Sec(NCA)^ is the most damaged species. We identified a 5′-tRNA fragment cleaved at the position 34 of tRNA^Sec(NCA)^ to promote the release of inflammatory cytokines and triggers innate immune response *via* activating the single-stranded RNA sensor, Toll-like receptor 7 (TLR7), which induces cellular senescence. Moreover, an antisense oligonucleotide targeting this fragment was developed to exhibit dual anti-aging effects, resulting in extending the healthspan and lifespan of naturally aged mice. These findings reveal tRNA damage as a novel aging hallmark and 5′-tRNA^Sec(NCA)^ half as a promising target for aging interventions.

## Results

### tRNA damage occurs in kidney during aging process

Kidney function progressively declines with age, leading to reduced glomerular filtration rate (GFR), impaired urine concentration, and decreased waste clearance, which increases the risk of chronic kidney disease, hypertension, and other metabolic disorders, ultimately impacting overall health and quality of life in the elderly^26^. Our group has previously developed a platform for characterization of tRNA species using specific nuclease digestion coupled with LC-MS analysis^27,28^. Thus, kidney of the adult (6-month-old) and aged (20-month-old) mice were harvested for comprehensively profiling of tRNAs, as well as protein expressions which would be impacted by any changes of tRNAs in aging (Figure 1a). Significant age-related morphological changes and reflecting glomerulosclerosis were observed in the kidneys by H&E staining (Supplementary Figure S1a). The results of tRNA mapping showed significant dysregulation in various tRNA species during aging, particularly a 70% decrease in tRNA^Sec(NCA)^ (Figure 1b). tRNA^Sec(NCA)^ is a unique tRNA that transfers its cognate amino acid selenocysteine to ribosomes for selenoprotein translation^29^. Although selenoproteins have been revealed to be associated with aging, the exact mechanisms remain unknown. To investigate the impact of tRNA damage on aging, we further analyzed protein levels in the kidneys of adult and aged mice using quantitative proteomics based on DIA-mode LC-MS technique, demonstrating that significant cellular pathway including immune system process and immune response were enriched in GO-BP enrichment (Figure 1c). Remarkably, several selenoproteins including TXNRD1, selenoprotein O (SelO), selenoprotein H (SelH), and GPX1 were downregulated in aged mice (Figure 1d), which was in consistent with the trend of their mRNA levels (Figure 1d, Supplementary Figure S1b). TXNRD1 exhibited a 64% decrease in aged mice, consistent with findings that TXNRD1 is downregulated in aged ovarian tissues compared to young mice^30^. Deficiency of SelO leads to vascular damage^31^, which exhibited a 27% decrease in aged mice. SelH, another thioredoxin-like nuclear protein with antioxidant and anti-aging effects, was observed with a 32% dysregulation in aged mice^32^. Together with the finding that GPX1 dysregulated for almost 49%, which catalyzes the reduction of hydrogen peroxide and organic hydroperoxides^33^, it suggests that tRNA^Sec(NCA)^ exhibits the most serious damage in aged mice, which leads to the downregulation of selenoproteins with anti-aging effects, potentially contributing to senescence in mice.

**Figure 1.**
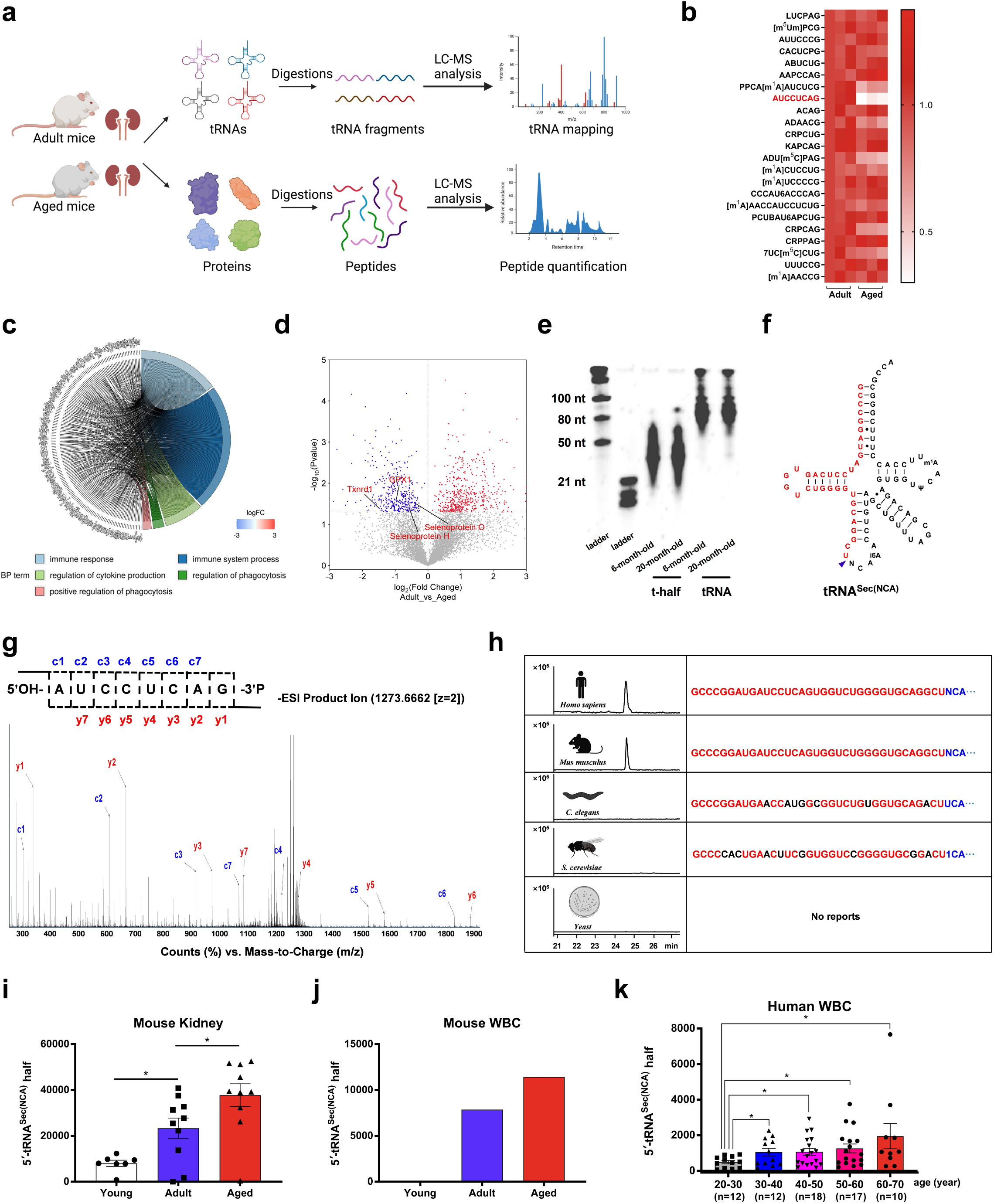
tRNA damage leads to 5′-tRNA^Sec(NCA)^ half accumulation during aging. **a**, Schematic workflow of the experiment demonstrating the process of extracting tRNAs from the kidneys of adult and aged mice, followed by digestion with RNase T1 and LC-MS analysis. Proteins were extracted for proteomic analysis. **b**, Heatmap of tRNA mapping in the kidneys of aged mice compared with adult mice. **c**, Proteomic analysis of the kidneys of adult and aged mice revealed significant enrichment of pathways such as immune system process and immune response in GO-BP enrichment. **d**, Volcano plot of selenoproteins level in the kidney from adult and aged mice. **e**, Enrichment of tRNA half fraction from the kidney of adult and aged mice. **f**, By comparing the theoretical MS data with the measured data, the RNA sequence corresponding to the ion m/z 2271.4690 was identified as the 5′-tRNA half cleaved at position 34 of tRNA^Sec(NCA)^. **g**, The purified RNA was further characterized as 5′-tRNA^Sec(NCA)^ half by RNase T1 digestion and LC-MS/MS analysis. **h**, 5′-tRNA^Sec(NCA)^ half was particularly detected in humans and mice, but not in *C. elegans*, *D. melanogaster*, and yeast. **i-j**, The 5′-tRNA^Sec(NCA)^ half was significantly accumulated in the kidneys and white blood cells of mice during aging. **k**, The abundance of 5′-tRNA^Sec(NCA)^ half increased with age in healthy human white blood cell samples. Data presented in **i** and **k** are presented as mean ± S.E.M. Statistical test: two-tailed unpaired *t*-test. \**P*<0.05; \*\**P*<0.01; \*\*\**P*<0.001; \*\*\*\**P*<0.0001; n.s., non-significant.

### tRNA half fraction from aged mice triggers higher innate immune responses

The aging process involves increasing physiological and psychological stress due to accumulated cellular damage, weakened immune function, and heightened disease susceptibility, impacting overall health and quality of life, which leads to tRNA damage to produce tRNA halves^25^. It has been addressed that small RNA fractions separated from the kidneys of 20-month-old rats could activate TLR7, which is a single-stranded RNA sensor, thus releasing inflammatory cytokines to induce aging^34^. This is consistent with our observation that aging markers including P16, P21, IL6, and CCL2, along with TLR7, were also upregulated in the kidney of aged mice (Supplementary Figure S1c). Thus, we speculated that tRNA halves induced by tRNA damage might induce aging *via* triggering innate immune responses. Transfection of tRNA half fraction isolated from the kidney of aged mice demonstrated a significantly higher TLR7 level compared to that of adult mice on macrophage RAW264.7 cells, which is a sensitive cell line commonly used in immunology (Supplementary Figure S1d). As a significant marker of immunomodulation, nitric oxide release of RAW264.7 cells was further detected^35^. The results indicated that the tRNA half fraction from aged mice exhibits significantly higher nitric oxide release than that from adult mice (Supplementary Figure S1e). These results suggest a variation in the biological functions of tRNA half fraction during aging.

### 5′-tRNASec(NCA) half is overexpressed during aging

Since tRNA^Sec(NCA)^ exhibited the most serious damage in kidneys of aged mice, we further aimed to assess the expression level of tRNA^Sec(NCA)^ half during aging. However, the cleavage site on tRNA molecule depends on complex ribonucleases that are difficult to predict. Therefore, tRNA half were artificially designed at different cleavage site of tRNA^Sec(NCA)^ sequence from MODOMICS database^36^. By comparing the theoretical MS data with the measured data collected from LC-MS profiling of adult and aged mice, an RNA sequence with the ion *m/z* 2271.4690 was matched as the -5 charge state of 5′-tRNA half cleaved at 34^th^ position of tRNA^Sec(NCA)^ (Figure 1f). Two-dimensional liquid chromatographic method integrating ion-pair and weak anion-exchange chromatographies was further purified from the kidney of aged mice (Supplementary Figure S2a)^37^. Furthermore, by using RNase T1 digestion coupled with LC-MS/MS analysis, the purified RNA was characterized as 5′-tRNA half cleaved at 34^th^ position of tRNA^Sec(NCA)^ (Figure 1g, Supplementary S2b, c, Supplementary Table S1), which was revealed to be significantly accumulated during aging process in mice kidney, heart and brain, as well as white blood cells (WBCs) (Figure 1h and i, Supplementary Figure S3a). Notably, the sequence information of the 5′-tRNA^Sec(NCA)^ half in mice and humans, as reported in the MODOMICS database, is identical^36^. Thus, we collected and analyzed the abundance of 5′-tRNA^Sec(NCA)^ half in 69 clinical WBCs samples from healthy human. The results indicate a progressive increase of 5′-tRNA^Sec(NCA)^ half in human WBCs with age, with statistically significant differences observed between the youngest age group (20-30 years old) and the older age groups (60-70 years old), suggesting an age-dependent alteration in tRNA^Sec(NCA)^ stability (Figure 1j). In addition, 5′-tRNA^Sec(NCA)^ half was undetected in *Caenorhabditis elegans*, *Drosophila melanogaster*, and yeast (*Saccharomyces cerevisiae*), which were used as negative controls due to the differing sequence information of tRNA^Sec(NCA)^ in these species compared to humans and mice (Figure 1k). These findings suggest that 5′-tRNA^Sec(NCA)^ half might play a role in aging process of both humans and mice, which can be served as a potential biomarker of aging.

ANG is usually responsible for tRNA cleavage at the anticodon loop to produce tRNA half^38^. However, qPCR analysis indicated an unexpected result that ANG levels in the kidney from aged mice were significantly lower than that from adult mice, which is opposite with the above findings that 5′-tRNA^Sec(NCA)^ half accumulates in aging process. Furthermore, both Dicer and RNase P upregulated in aged mice (Supplementary Figure S3b). These results demonstrated that the cleavage of tRNA^Sec(NCA)^ to produce tRNA half might do not follow the classic pathway of tRNA cleavage.

### 5′-tRNASec(NCA) half induces cellular senescence phenotype

Since 5′-tRNA^Sec(NCA)^ half exhibited the most accumulation in aging process, we speculated that it is a major player of tRNA half fraction to induce senescence. Therefore, we explored the direct effects of synthetic 5′-tRNA^Sec(NCA)^ half on cellular senescence in mouse renal tubular epithelial cells (TCMK-1) and human diploid fibroblast cells (2BS). By employing liposomal transfection, CCK-8 detection showed that the proliferation of both cell lines treated with 5′-tRNA^Sec(NCA)^ half was significantly inhibited compared to the blank transfection reagents group (Figure 2a). SACβCgal staining demonstrated that both cells treated with 5′-tRNA^Sec(NCA)^ half exhibited a significantly increased SACβCgal-positive rate, suggesting the phenotype of cellular senescence (Figure 2b). In consistent, aging markers including P21, P16, IL6, and CCL2 in both cells treated with 5′-tRNA^Sec(NCA)^ half significantly upregulated (Figure 2c). Furthermore, reactive oxygen species (ROS) as a phenotype of inflammation significantly increased in both cells treated with 5′-tRNA^Sec(NCA)^ half (Figure 2d). As another senescence marker, telomere length influenced by telomerase activity shortens with aging. To further address aging-inducing ability of 5′-tRNA^Sec(NCA)^ half, the T/S ratio as a quantitative measure obtained by comparing the amount of telomere repeat copy number (T) to a single-copy gene (S) was analyzed. The results demonstrated that telomere length significantly decreased in both 5′-tRNA^Sec(NCA)^ half-treated cells (Figure 2e). Furthermore, IFN-β levels significantly increased upon treatment with 5′-tRNA^Sec(NCA)^ half, while IFN-γ levels remained unchanged (Figure 2f). This phenomenon was accordingly verified in human plasma that the IFN-β levels were significantly upregulated from 40-to 50-year-old people to that in 60- to 70-year-old people (Supplementary Figure S4a). Accordingly, bioinformatics analysis of genome data from TCGA database addressed that, compared to paracancerous tissue, both TLR7 and IFNB1 (encoding IFN-β) exhibits high expressions in renal tumors that might be an outcome of kidney aging (Supplementary Figure S4b, c). These findings indicate that 5′-tRNA^Sec(NCA)^ half might induce senescence *via* activating type-I interferon to release inflammatory cytokines.

**Figure 2.**
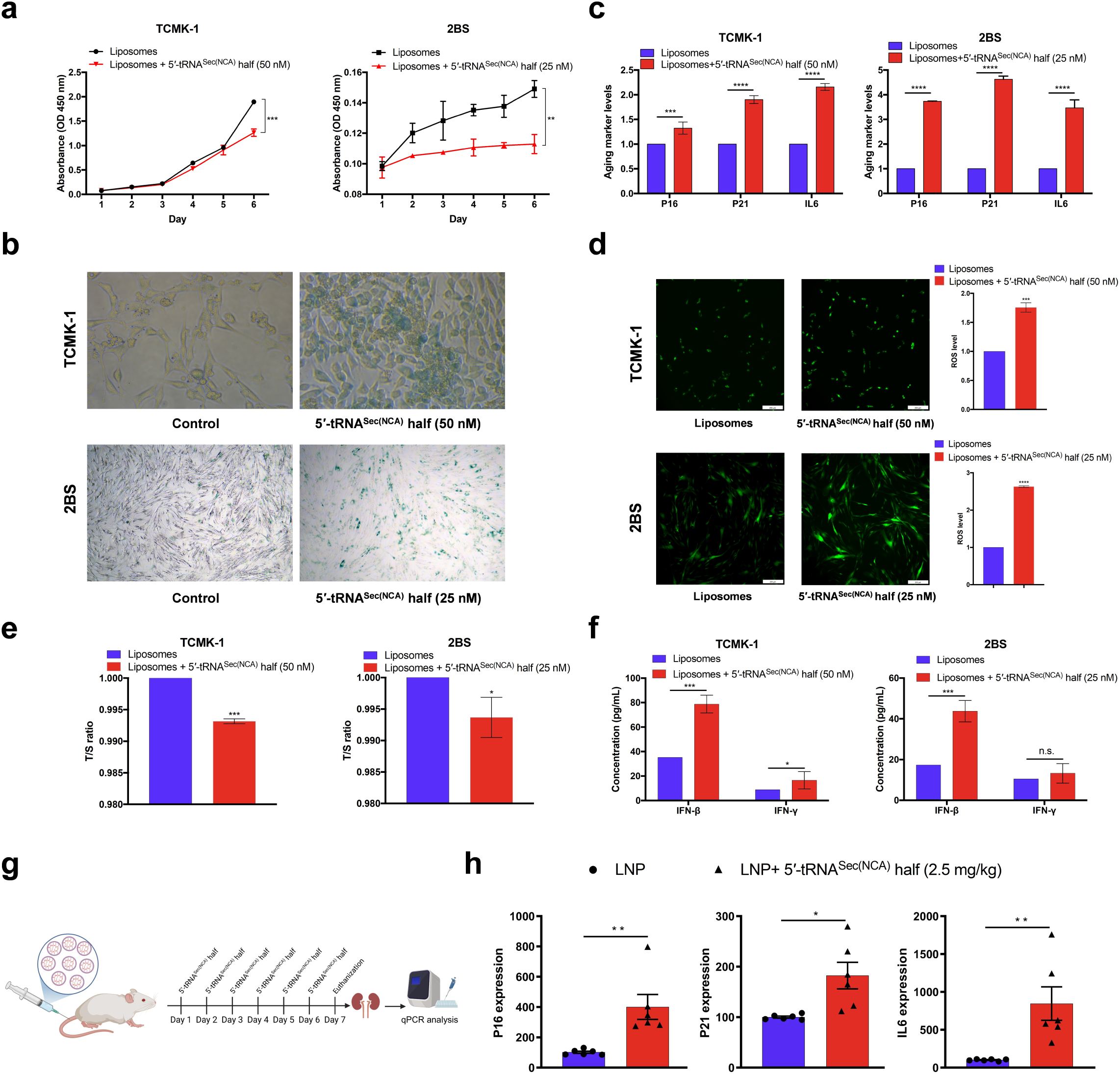
5′-tRNA^Sec(NCA)^ half induces cellular senescence phenotype. **a**, CCK-8 assay showed that the proliferation of TCMK-1 and 2BS treated with 5′-tRNA^Sec(NCA)^ half was significantly inhibited compared with the blank transfection reagent group. **b**. SA-β-gal staining indicated that the SA-β-gal positive rate of TCMK-1 and 2BS cells treated with 5′-tRNA^Sec(NCA)^ half increased significantly, presenting a cellular senescence phenotype. **c**, The expression levels of senescence markers P21, P16, IL6, and CCL2 were significantly upregulated in cells induced by 5′-tRNA^Sec(NCA)^ half. **d**, ROS significantly increased in cells treated with 5′-tRNA^Sec(NCA)^ half. **e**, The telomere length (T/S ratio) of cells treated with 5′-tRNA^Sec(NCA)^ half significantly decreased. **f**. IFN-β levels significantly upregulated in cells treated with 5′-tRNA^Sec(NCA)^ half, while the level of IFN-γ did not exhibit such huge change. **g-h**, Liposome nanoparticle (LNP) delivery system encapsulated with 5′-tRNA^Sec(NCA)^ half was intravenously injected into young Balb/c female mice. After 6 days, the senescence markers in the kidneys of mice were significantly upregulated. Data presented in **c**, **d**, **e**, and **f** are presented as mean ± S.D. Data presented in **h** are presented as mean ± S.E.M. Statistical test: two-tailed unpaired *t*-test. \**P*<0.05; \*\**P*<0.01; \*\*\**P*<0.001; \*\*\*\**P*<0.0001; n.s., non-significant.

To further confirm the above findings, liposome nanoparticle (LNP) delivery systems was applied to encapsulate 5′-tRNA^Sec(NCA)^ half and intravenously injected into young Balb/c female mice (1-month-old) with the dose of 2.5 mg/kg/day. After constantly administration for 6 days, mice were sacrificed and aging markers were found to be significantly upregulated in the kidney of mice treated with 5′-tRNA^Sec(NCA)^ half (Figure 2g, h). The above results suggest that 5′-tRNA^Sec(NCA)^ half induces senescence both *in vitro* and *in vivo*.

### 5′-tRNASec(NCA) half activates TLR7 to trigger innate immune response

Previous studies have revealed 5′-half molecules of tRNA^His^(GUG) as an abundant activator of TLR7 in human monocyte-derived macrophages-secreted extracellular vehicles, as well as 5′-tRNA^Val^(CAC)^/^(AAC) half activating TLR7 in the plasma of patients infected with *Mycobacterium tuberculosis*^39,40^. Thus, we speculated that the increase of type I interferon releasing induced by 5′-tRNA^Sec(NCA)^ half is due to the activation of TLR7. Such activation would lead to the upregulation of other RNA sensor genes *via* the positive-feedback loop of the innate immune responses^41^. Further transcriptomic sequencing revealed that RNA sensor genes, including TLR7, MDA5, RIG-I, TLR3, OAS2, and OAS3, were upregulated in 5′-tRNA^Sec(NCA)^ half-treated cells (Figure 3a). This finding was corroborated by qPCR analysis, which showed significant increase in the expression of the above genes, including MYD88, a critical adaptor protein that plays a key role in the signal transduction pathway of TLR7 (Figure 3b). In consistent, the protein level of TLR7 in TCMK-1 and 2BS cells treated with 5′-tRNA^Sec(NCA)^ half significantly upregulated in a dose-dependent manner (Figure 3c).

**Figure 3.**
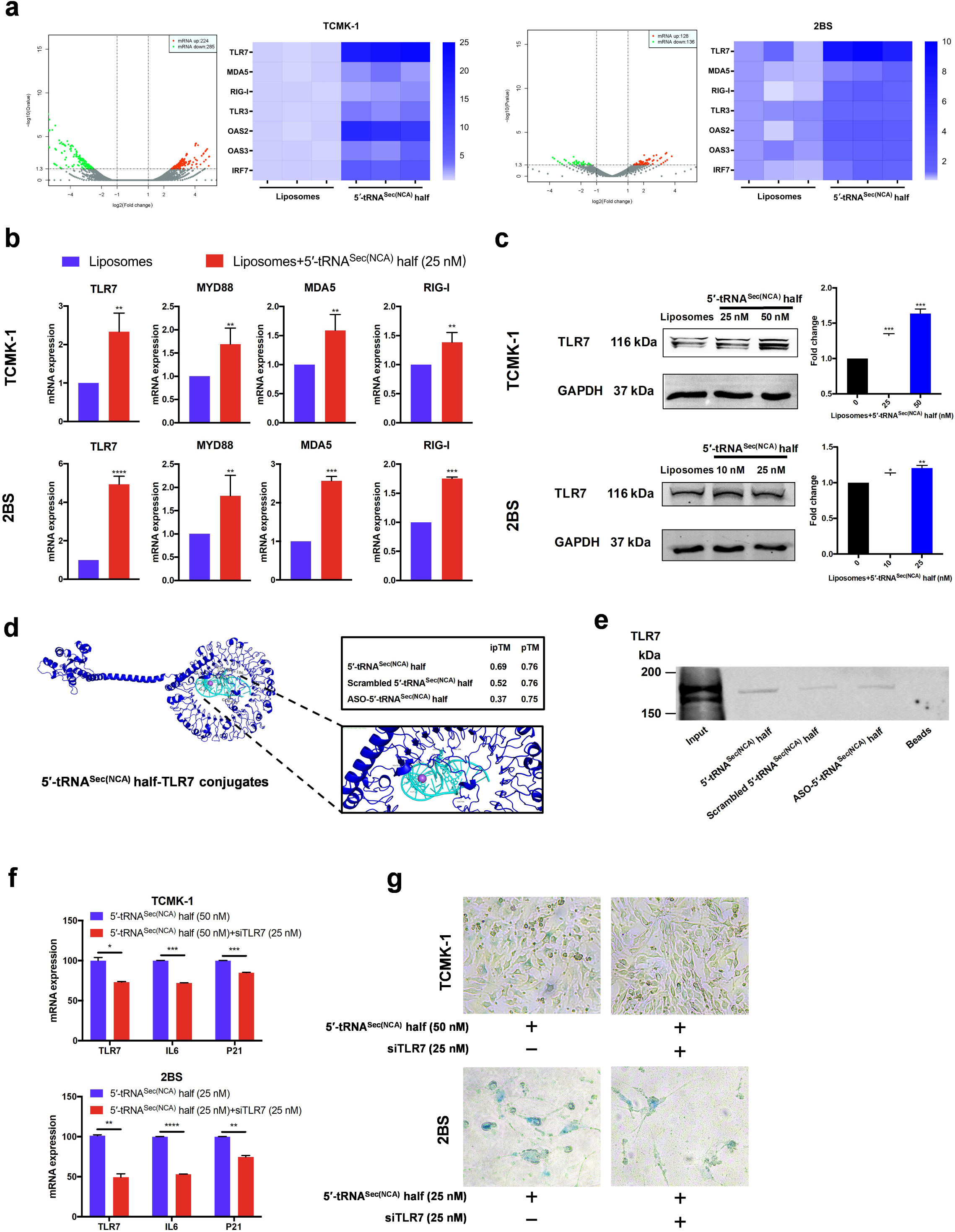
5′-tRNA^Sec(NCA)^ half activates TLR7 to trigger innate immune response. **a**, Transcriptomic sequencing revealed that dsRNA sensor genes significantly upregulated in cells treated with 5′-tRNA^Sec(NCA)^ half. **b**, qPCR analysis confirmed that the expression of the above genes and MYD88, a key adaptor protein in the TLR7 signaling pathway, increased significantly. **c**, In TCMK-1 and 2BS cells treated with 5′-tRNA^Sec(NCA)^ half, the protein level of TLR7 was significantly upregulated in a dose-dependent manner. **d**, Simulation of the interaction between 5′-tRNA^Sec(NCA)^ half and TLR7 protein predicted by AlphaFold 3. **e**, RNA pull-down assay demonstrated that 5′-tRNA^Sec(NCA)^ half specifically bound to TLR7. **f-g**, In TLR7-knockdown (TLR7-KD) cells, co-treatment with 5′-tRNA^Sec(NCA)^ half and siTLR7 led to a significant downregulation of senescence markers and a lower SA-β-gal positive rate compared with treatment with 5′-tRNA^Sec(NCA)^ ^half^ alone. Data presented in **b**, **c**, and **f** are presented as mean ± S.D. Statistical test: two-tailed unpaired *t*-test. \**P*<0.05; \*\**P*<0.01; \*\*\**P*<0.001; \*\*\*\**P*<0.0001.

We further use the accessible AlphaFold 3 Server of AF3 to predict the interaction of 5′-tRNA^Sec(NCA)^ half and TLR7. The results demonstrated that 5′-tRNA^Sec(NCA)^ half is more possible to interact with TLR7 rather than its scramble and antisense sequence (Figure 3d). An RNA pull-down assay was further performed to investigate the potential interaction, showing a distinct protein band corresponding to TLR7 in 5′-tRNA^Sec(NCA)^ half-treated proteins isolated from kidney cells, whereas negative RNA-treated samples (scrambled sequence and antisense oligonucleotides) exhibited much less lower expression in this band (Figure 3e). These results indicate a sequence-specific binding of 5′-tRNA^Sec(NCA)^ half to TLR7. Functionally, we validated the effect of 5′-tRNA^Sec(NCA)^ half in TLR7-knockdown (TLR7-KD) cells treated with siRNA. The results indicated a significant downregulation of aging markers (P21 and IL6), as well as SA-β-gal-positive rate in cells co-treated by 5′-tRNA^Sec(NCA)^ half and siTLR7 than 5′-tRNA^Sec(NCA)^ half used alone (Figure 3f, g). Collectively, these results indicate that 5′-tRNA^Sec(NCA)^ half specifically binds to TLR7 protein to induce cellular senescence.

### ASO-5**′**-tRNA^Sec(NCA)^ half intervention ameliorates cellular senescence

Since 5′-tRNA^Sec(NCA)^ half was suggested to induce senescence *in vitro* and *in vivo*, we selected antisense oligonucleotide (ASO) of 5′-tRNA^Sec(NCA)^ half to investigate its potential intervention effect on aged cell models, including D-galactose-induced TCMK-1 aged cells and 2BS naturally aged cells (passage 37), in which 5′-tRNA^Sec(NCA)^ half levels were significantly higher than normal cells (Figure 4a). Initially, a CCK-8 assay addressed that the proliferation of both aged cells treated with ASO-5′-tRNA^Sec(NCA)^ half significantly increased compared to liposome-treated controls (Figure 4b). Consistently, ASO-5′-tRNA^Sec(NCA)^ half treatment resulted in a significant decrease in SA-β-gal-positive cells in both aged cells (Figure 4c). Additionally, ROS levels were markedly reduced in ASO-5′-tRNA^Sec(NCA)^ half-treated aged cells (Figure 4d). Furthermore, ASO-5′-tRNA^Sec(NCA)^ half significantly downregulated aging markers and extended telomere length in both aged cells (Figure 4e, f).

**Figure 4.**
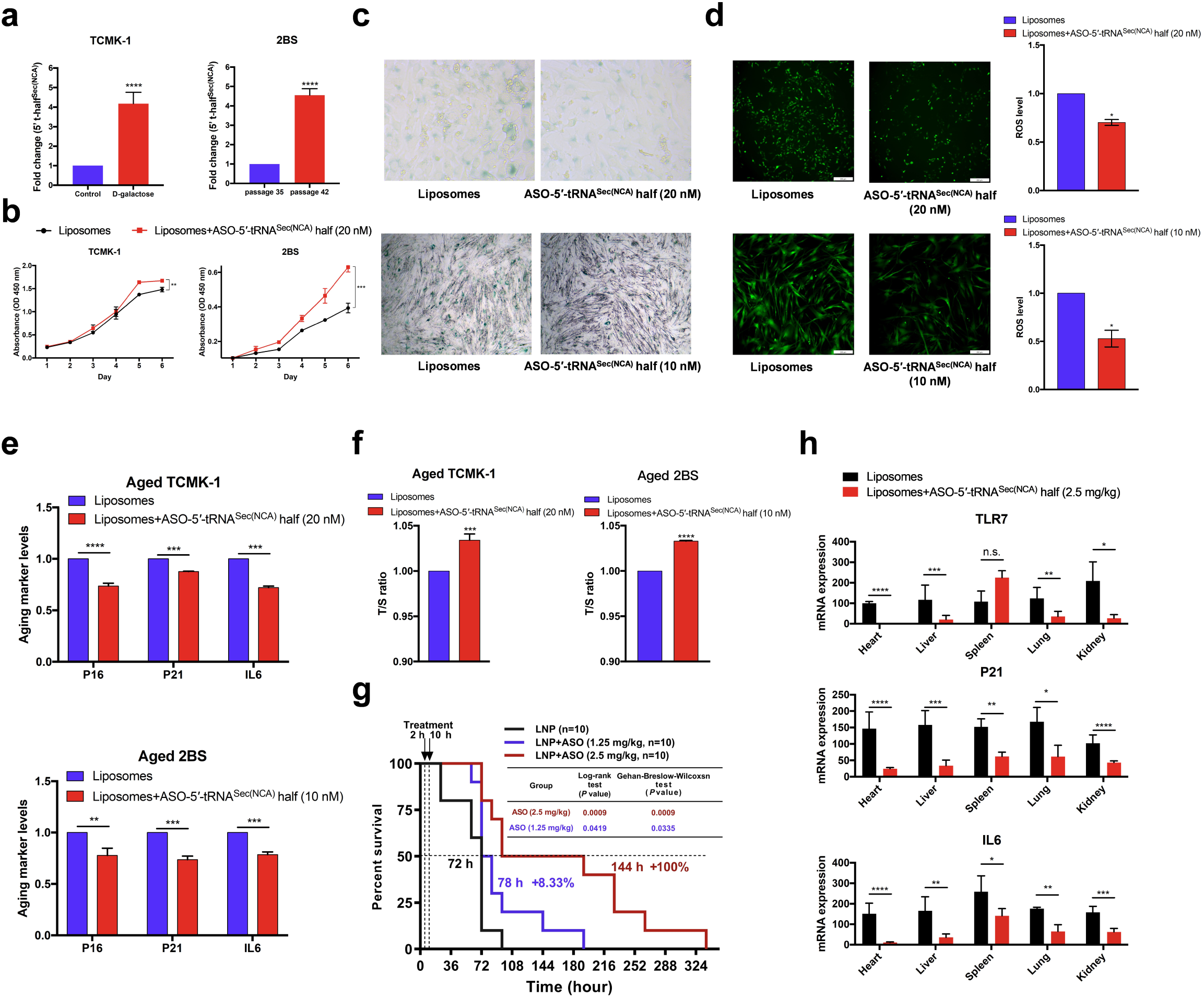
ASO-5′-tRNA^Sec(NCA)^ half intervention ameliorates cellular senescence. **a**, 5′-tRNA^Sec(NCA)^ half significantly increased in D-galactose-induced TCMK-1 aged cells and 2BS naturally aged cells. **b**, CCK-8 assay indicated that the proliferation of aged cells treated with ASO-5′-tRNA^Sec(NCA)^ half significantly increased compared with the liposome-treated controls. **c**. Treatment with ASO-5′-tRNA^Sec(NCA)^ half resulted in a significant decrease in the number of SA-β-gal-positive cells in aged cells. **d**, ROS levels were markedly reduced in aged cells treated with ASO-5′-tRNA^Sec(NCA)^ half. **e-f**. ASO-5′-tRNA^Sec(NCA)^ half significantly downregulated aging markers and extended telomere length in aged cells. **g**, LNP delivery of ASO-5′-tRNA^Sec(NCA)^ half significantly extended the lifespan of paraquat-induced acute aged mice in a dose-dependent manner, as well as dysregulating TLR7, P21 and IL6 in multiple organs (**h**). Data presented in **a**, **b**, **d**, **e**, **f** and **h** are presented as mean ± S.D. Data presented in **g** are presented as log-rank test and the Gehan-Breslow-Wilcoxon. Statistical test: two-tailed unpaired *t*-test. \**P*<0.05; \*\**P*<0.01; \*\*\**P*<0.001; \*\*\*\**P*<0.0001; n.s., non-significant.

To evaluate the potential anti-aging effects of ASO-5′-tRNA^Sec(NCA)^ half *in vivo*, a paraquat-induced acute aging model was employed in Balb/c female mice. ASO-5′-tRNA^Sec(NCA)^ half encapsulated in LNP was intravenously injected at 0 and 8 h after paraquat administration. Remarkably, compared to the LNP control group, ASO-5′-tRNA^Sec(NCA)^ half dose-dependently extended lifespan (as assessed at median lifespan) in mice by 100% of 2.5 mg/kg dose (*P* = 0.0009 by both log-rank test and Gehan-Breslow-Wilcoxon test) and 8.33% of 1.25 mg/kg dose (*P* = 0.0419 by log-rank test and *P* = 0.0335 by Gehan-Breslow-Wilcoxon test), respectively (Figure 4g). In consistent, significant reduction of aging markers (P16, P21, IL6) and TLR7 was observed in heart, liver, spleen, lung, and kidney, compared to the LNP-treated group (Figure 4h).

Inspired by the acute aging model *in vivo*, naturally aged mice were further employed to evaluate the anti-aging effects of ASO-5′-tRNA^Sec(NCA)^ half in whole lifespan. Therefore, 16-month-old male C57 mice were intravenously injected with LNP-encapsulated ASO-5′-tRNA^Sec(NCA)^ half twice a month till 25-month-old (treated group), while blank LNP treatment was used as control group (Figure 5a). Meanwhile, for female mice, 20-month-old Balb/c mice were intravenously injected with LNP-encapsulated ASO-5′-tRNA^Sec(NCA)^ half twice a month till 29-month-old (Supplementary Figure S5a). The fur of male mice in the 20-month-old control group presented remarkable whitening and substantial hair loss. By contrast, male mice that received ASO-5′-tRNA^Sec(NCA)^ half treatment demonstrated a relatively lower level of hair loss when compared to the LNP control group (Figure 5b). Remarkably, compared to the LNP control group, ASO significantly extended lifespan (as assessed at median lifespan) in male and female mice by 24.1% and 12.4%, respectively (*P* <0.0001 by log-rank test and *P* = 0.0003 by Gehan-Breslow-Wilcoxon test, Figure 5c and Supplementary Figure S5b). Obviously, ASO-5′-tRNA^Sec(NCA)^ half-treated mice exhibited a 5.7% reduction in age-associated body weight gain and less alopecia compared to the LNP-treated group at 25-month-old, while female mice exhibited a 7.6% reduction at 21.5-month-old (Figure 5d, Supplementary Figure S5c). 3D X-ray imaging indicated significantly higher bone mineral density in ASO-5′-tRNA^Sec(NCA)^ half-treated mice of both sexes, which have significant higher lean percent and lower fatty percent (Figure 5e-f, Supplementary Figure S5d). In an open-field test, ASO-5′-tRNA^Sec(NCA)^ half-treated mice of both sexes demonstrated significantly increased movement (Figure 5h, Supplementary Figure S5e). The Y maze test showed that ASO-5′-tRNA^Sec(NCA)^ half-treated mice of both sexes had significantly higher alteration rates, indicating enhanced exploratory behavior (Figure 5i, Supplementary Figure S5f). Grip strength tests revealed that ASO-5′-tRNA^Sec(NCA)^ half-treated mice of both sexes had higher grip strength (Figure 5j, Supplementary Figure S5g). In the rotarod test, ASO-5′-tRNA^Sec(NCA)^ half-treated mice of both sexes displayed significantly longer hanging times and greater distances traveled compared to control mice (Figure 5k, Supplementary Figure S5h). Notably, there is no significant difference of the immobility in mice of both sexes treated by ASO-5′-tRNA^Sec(NCA)^ half, suggesting ASO-5′-tRNA^Sec(NCA)^ half did not induce anxiety (Figure 5l, Supplementary Figure S5i).

**Figure 5.**
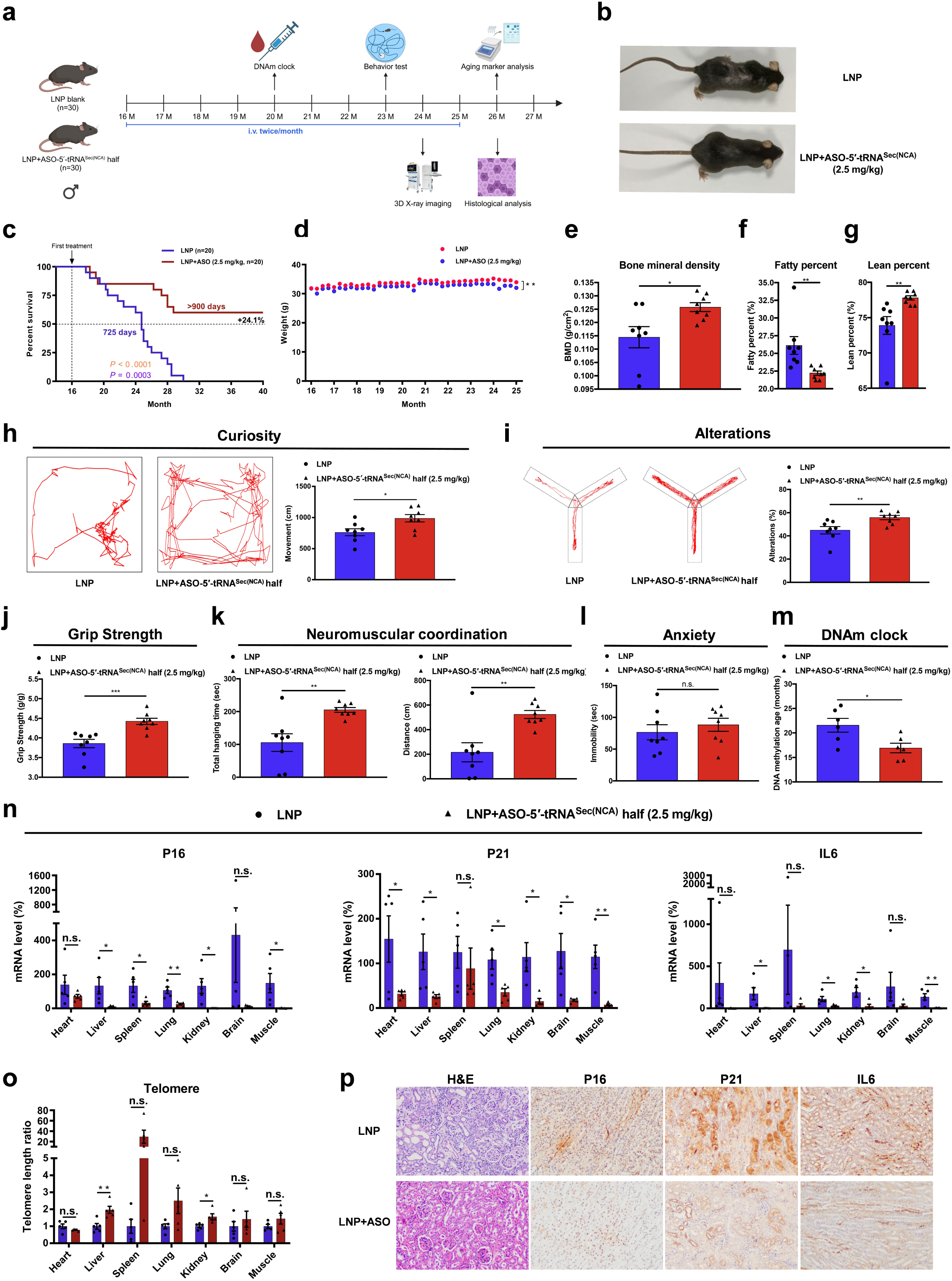
ASO-5′-tRNA^Sec(NCA)^ half extends the healthy lifespan of naturally aged mice. **a**, Experimental design of anti-aging test of ASO-5′-tRNA^Sec(NCA)^ half from 16-month-old mice. **b**, Image of 20-month-old mice received LNP encapsulated ASO-5′-tRNA^Sec(NCA)^ half treatment or not. Survival curves (**c**) and weight curve (**d**) of aged mice received LNP encapsulated ASO-5′-tRNA^Sec(NCA)^ half treatment (n=20) or not (n=20). **e**, Bone mineral density measured by 3D X-ray imaging. **f-g**, Mice treated with ASO-5′-tRNA^Sec(NCA)^ half had a significantly higher lean percent and a lower fatty percent. **h**, Movement of mice received ASO-5′-tRNA^Sec(NCA)^ half treatment or not in the open-field test. **i**, Alteration rates of ASO-5′-tRNA^Sec(NCA)^ half-treated mice or not in the Y maze test. Grip strength test (**j**) and rotarod test (**k**) of ASO-5′-tRNA^Sec(NCA)^ half-treated mice or not. **l**, There was no significant difference in the immobility time between the two groups of mice. **m**, RRBS sequencing of DNA methylation age showed that 20-month-old aged mice treated with ASO-5′-tRNA^Sec(NCA)^ half were approximately 18.6 weeks younger than the LNP control group. Aging markers expressions (**n**) and telomere length (**o**) from the major organs of ASO-5′-tRNA^Sec(NCA)^ half-treated mice or not. **p**, H&E staining and IHC staining of P16, P21, and IL6 in major organs from mice treated with ASO-5′-tRNA^Sec(NCA)^ half or not. Data presented in **d**, **e**, **f**, **g**, **h**, **i**, **j**, **k**, **l**, **m**, **n**, and **o** are presented as mean ± S.E.M. Statistical test: two-tailed unpaired *t*-test. \**P*<0.05; \*\**P*<0.01; \*\*\**P*<0.001; n.s., non-significant. Data presented in **c** are presented as the log-rank test in orange and the Gehan-Breslow-Wilcoxon in purple.

DNA methylation age has widely applied to demonstrate the biological age of independent livings^15,42^. The multi-tissue predictor of DNA methylation age developed was applied to evaluate the anti-aging effect of ASO-5′-tRNA^Sec(NCA)^ half in natural aged male mice. As shown in Figure 5m, the results showed that aged male mice treated with ASO-5′-tRNA^Sec(NCA)^ half were approximately 18.6 weeks younger than the LNP control group (LNP+ASO-5′-tRNA^Sec(NCA)^ half=67.66 DNA-methylation-weeks [SD, 8.75]; LNP=86.27 DNA-methylation-weeks [SD, 12.64]). Furthermore, a significant downregulation of IL-6 was observed in the plasma of ASO-5′-tRNA^Sec(NCA)^ half-treated male mice (Supplementary Figure S6a). Aging markers in WBCs indicated that ASO-5′-tRNA^Sec(NCA)^ half-treated male mice exhibited significantly lower expressions, while not in the expression of telomerase (Supplementary Figure S6b, c). Complete blood analysis indicated that male mice treated with ASO-5′-tRNA^Sec(NCA)^ half of both sexes have significant higher level of PLT, PCT, MCH, MCV, HCT, HGB, RBC, EOS, and BAS (Supplementary Figure S6d). Furthermore, biochemical indicators TG, CRE, TC, and LDL-c in plasma significantly dysregulated in ASO-5′-tRNA^Sec(NCA)^ half-treated male mice, while there were no significant differences in other indicators such as glucose, HDL-c, AST, GGT, and ALT (Supplementary Figure S6e).

The trend of complete blood analysis of female mice is in accordance to that of male mice, as well as their biochemical indicators except for GGT which dysregulated in ASO-5′-tRNA^Sec(NCA)^ half-treated female mice (Supplementary Figure S7a, b). Remarkably, female reproductive hormones such as PROG, AMH and E2 of female mice upregulated upon ASO-5′-tRNA^Sec(NCA)^ half treatment (Supplementary Figure S7c). The above results indicated that ASO treatment not only ameliorates kidney function and exhibits safety for liver in aged mice of both sexes, but also implies its potential to improve ovarian function, enhance reproductive capacity, and delay organismal aging.

Concurrently, most organs from ASO-5′-tRNA^Sec(NCA)^ half-treated male mice exhibited significantly lower expressions of P16, P21 and IL6 (Figure 5n). In consistent, telomere length of major organs from ASO-5′-tRNA^Sec(NCA)^ half-treated male mice were significant longer than that from LNP-treated mice (Figure 5o). H&E staining revealed that kidneys from male mice treated with ASO-5′-tRNA^Sec(NCA)^ exhibited significantly healthier morphology compared to those from LNP-treated mice. This was further supported by IHC staining, which showed markedly reduced expression of aging markers such as P16, P21, and IL6, as well as similar improvements in other major organs (Figure 5p, Supplementary Figure S8). In summary, ASO-5′-tRNA^Sec(NCA)^ half-treated mice of both sexes exhibited a significantly longer healthy lifespan compared to the LNP control group.

### ASO-5**′**-tRNA^Sec(NCA)^ half targets and downregulates BCAT1 mRNA to suppress aging

Transcriptomic sequencing was further carried out to investigate the potential impact of ASO-5′-tRNA^Sec(NCA)^ half in genes. Remarkable downregulation of interferon-associated genes was observed in ASO-5′-tRNA^Sec(NCA)^ half-treated aged TCMK-1 and 2BS cells (Figure 6a). Gene Ontology (GO) analysis revealed that a longevity-associated pathway was enriched in ASO-5′-tRNA^Sec(NCA)^ half-treated aged cells, highlighting the significant downregulation of BCAT1, which dysregulated during aging (Figure 6b). Inhibition of BCAT1 *via* RNAi method has been shown to extend median lifespan by 25% in *C. elegans*^43^. Bioinformatics analysis predicted that ASO-5′-tRNA^Sec(NCA)^ half has a putative binding site in the 3′UTR region of BCAT1 mRNA. In *Mus musculus* (mice), the complementarity rate surpassed 75%, indicating a potential RNAi effect of ASO-5′-tRNA^Sec(NCA)^ half on BCAT1 dysregulation. However, in *Homo sapiens* (humans), the complementarity rate was only 45%, suggesting that the dysregulation of BCAT1 in humans is more likely to be attributed to other signaling pathways rather than the direct RNAi effect of ASO-5′-tRNA^Sec(NCA)^ half (Figure 6c). To investigate whether ASO-5′-tRNA^Sec(NCA)^ half directly targets BCAT1 of mice, the dual-luciferase reporter assay was utilized. The luciferase reporter gene vectors of BCAT1-wild type (WT) and -mutant (MUT) were co-transfected into HEK-293T cells together with either the ASO-5′-tRNA^Sec(NCA)^ half or its scrambled sequence. The outcomes indicated that the ASO-5′-tRNA^Sec(NCA)^ half remarkably decreased the luciferase activity of the BCAT1-WT reporter vector, while no significant alteration in the luciferase activity of the BCAT1-MUT reporter gene vector was detected (Figure 6d). qPCR analysis confirmed that ASO-5′-tRNA^Sec(NCA)^ half significantly inhibits BCAT1 expression in aged cells (Figure 6e). In consistent, BCAT1 at mRNA and protein level was significantly downregulated in the kidneys of paraquat-induced mice treated with ASO-5′-tRNA^Sec(NCA)^ half (Figure 6f, g). In naturally aged mice, major organs including liver, spleen, lung, kidney, and muscle from mice receiving ASO-5′-tRNA^Sec(NCA)^ half treatment exhibited remarkable lower mRNA levels of BCAT1 (Figure 6h). Moreover, the dysregulation of BCAT1 was also confirmed in these organs by IHC staining (Figure 6i). Overall, these findings demonstrate that the ASO not only blocked senescence-induing effect of 5′-tRNA^Sec(NCA)^ half, but also functions through RNAi pathway to suppress BCAT1 mRNA, thereby extending the healthspan and lifespan of naturally aged mice (Figure 7).

**Figure 6.**
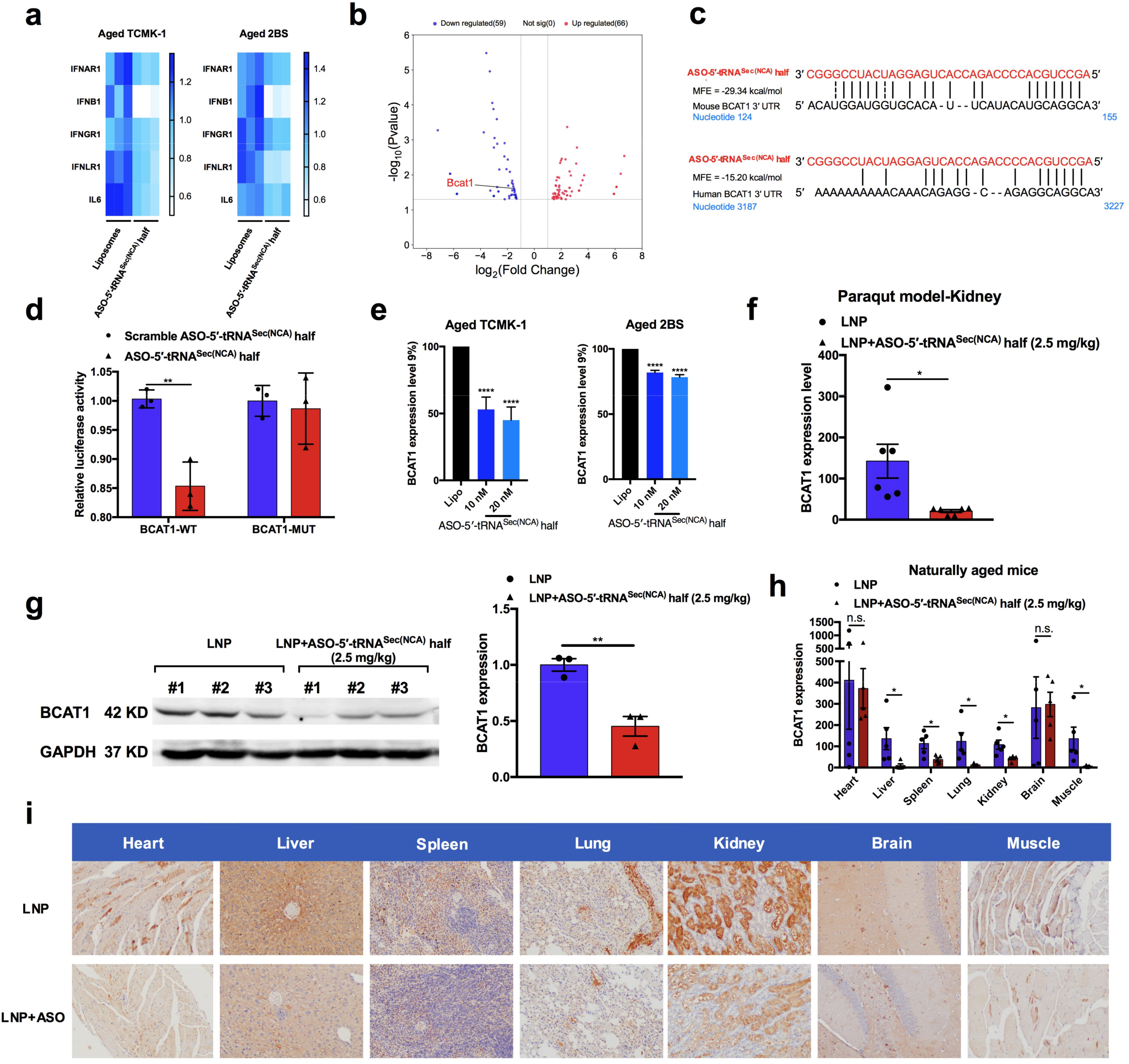
ASO-5′-tRNA^Sec(NCA)^ half targets and downregulates BCAT1 mRNA to suppress aging. **a**, Heatmap of interferon-associated genes in aged cells treated with ASO-5′-tRNA^Sec(NCA)^ half or not by transcriptomic sequencing. **b**, Volcano plot revealed that BCAT1 dysregulated in ASO-5′-tRNA^Sec(NCA)^ half-treated TCMK-1 cells. **c**, Putative binding sites of ASO-5′-tRNA^Sec(NCA)^ half on the 3′UTR region of BCAT1 mRNA of mice and humans. **d**, The dual-luciferase reporter assay demonstrated that ASO-5′-tRNA^Sec(NCA)^ half remarkably decreased the luciferase activity of mice BCAT1-WT reporter vector, while no significant alteration was detected in the luciferase activity of mice BCAT1-MUT reporter gene vector. **e**, qPCR analysis confirmed that ASO-5′-tRNA^Sec(NCA)^ half significantly inhibited BCAT1 expression in aged cells. **f-g**, In paraquat-induced mice treated with ASO-5′-tRNA^Sec(NCA)^ half, BCAT1 was downregulated at both the mRNA and protein levels in the kidneys. **h**, Major organs including liver, spleen, lung, kidney, and muscle from naturally aged mice receiving ASO-5′-tRNA^Sec(NCA)^ half treatment exhibited remarkably lower mRNA levels of BCAT1. **i**, IHC staining of BCAT1 in major organs from naturally aged mice. Data presented in **d** and **e** are presented as mean ± S.D. Data presented in **f**, **g**, and **h** are presented as mean ± S.E.M. Statistical test: two-tailed unpaired *t*-test. \**P*<0.05; \*\**P*<0.01; \*\*\*\**P*<0.0001; n.s., non-significant.

**Figure 7.**
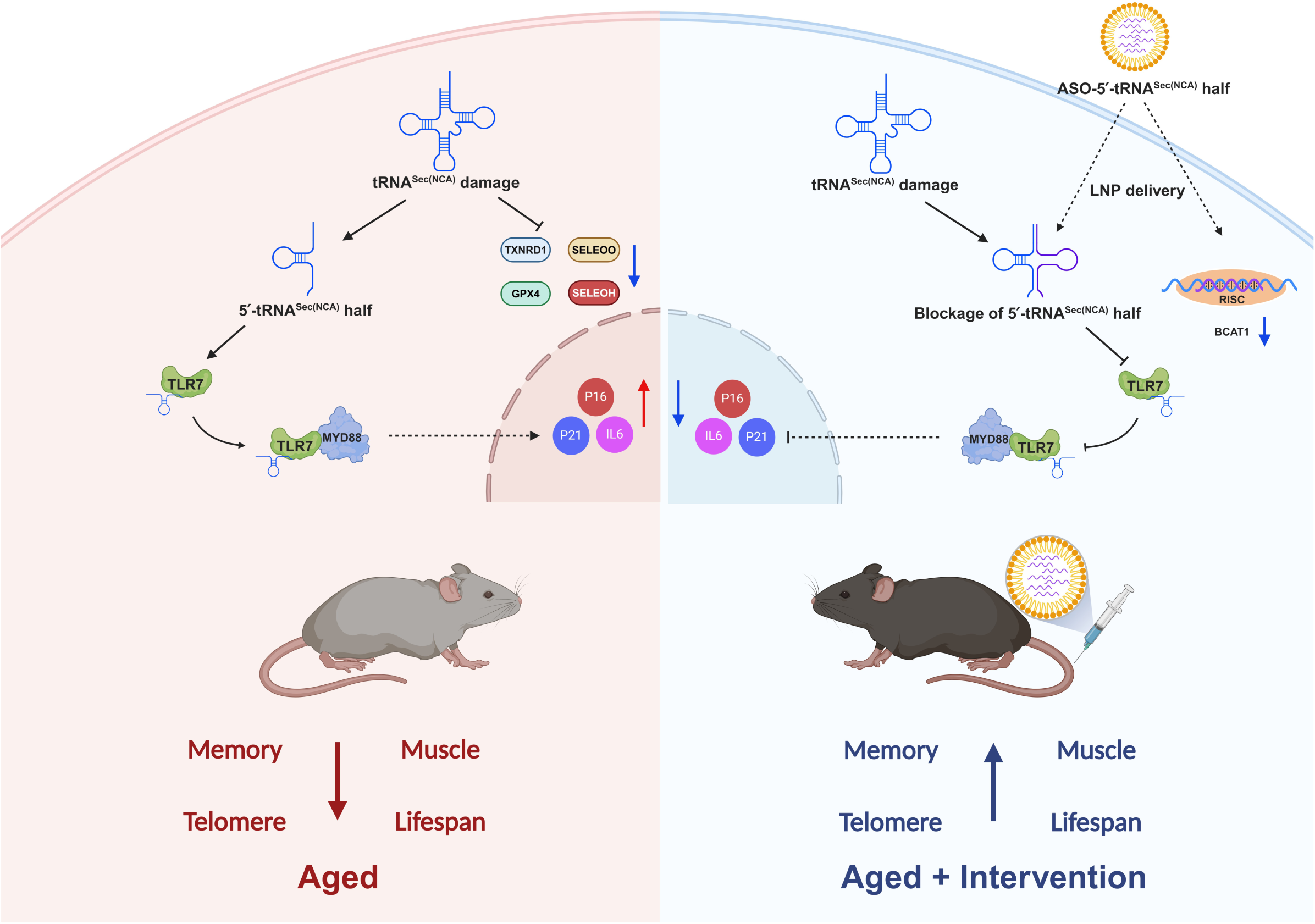
Working model of this study.

## Discussion

The aging process engenders a distinct stress-inducing cellular milieu characterized by progressive functional decline. While epigenetic alterations including DNA methylation shifts and histone modification alterations has been extensively characterized as a pillar of aging biology, the emerging role of non-coding RNAs (ncRNAs) in age-related pathways have also been extensively explored. Small extracellular vesicles containing miRNAs from the plasma of young mice stimulate PGC-1α expression through miRNA species^44^. In aged mice, an aging-related lncRNA specifically expressed in nucleus of regulatory T (T_reg_) cells causes T_reg_ metabolic dysfunction^45^. It was also identified that an insulin-sensitive circular RNA in Drosophila regulates lifespan through insulin/IGF-1 signaling pathway^46^. miR-188-3p targets skeletal endothelium to couple angiogenesis and osteogenesis during aging^47^. Meanwhile, miR-455-3p enhances synaptic and cognitive functions and extends lifespan^48^. These findings suggest that the epigenetic alterations are more likely to function as primary initiators of the aging process, such as tRNAs and their fragments. Notably, tRNA fragmentation primarily arises from cellular stress challenges. The kidney represents a particularly vulnerable organ in this context, being constantly exposed to high physiological stress from its blood filtration functions and metabolic waste processing. This inherent vulnerability is further exacerbated by pathological stressors including diabetes, hypertension, autoimmune disorders, and nephrotoxic drug exposure, collectively creating an exceptionally high-stress microenvironment conducive to tRNA damage^49^. Our study addresses that, as a direct player to regulate selenoproteins translation, tRNA^Sec(NCA)^ damage remarkedly leads the downregulation of selenoproteins with anti-aging effect. This, together with the finding that knocking out the tRNA^Sec^ encoded by Trsp substantially decreases selenoprotein content and results in Kashin-Beck disease^50^, demonstrates that selenoproteins dysregulation induces aging. Moreover, tRNA^Sec(NCA)^ damage produced 5′-tRNA^Sec(NCA)^ half, which accumulates during aging process in multiple organs of mice. This phenomenon is then extensively verified in human WBCs but not in other natural aging models such as *C. elegans*, *D. melanogaster* and yeast due to their difference in transcriptome, which emphasized the importance of 5′-tRNA^Sec(NCA)^ half as a significant marker of aging in both mice and humans. The observed dysregulation of tRNA^Sec(NCA)^ and its impact on the expression of selenoproteins are vital for cellular redox homeostasis, indicating the importance of tRNA integrity in maintaining cellular function during aging. Therefore, our findings suggest that tRNA damage should also be considered as a crucial epigenetic hallmark and is of great significance in the field of aging research, providing new directions for in-depth exploration of aging mechanisms.

Overexpressed innate immune responses lead to the release of pro-inflammation cytokines that induce cellular senescence, results in inflammaging^51^. Toll-like receptor (TLR) family plays a pivotal role in innate immune responses and has been implicated in the aging process^52^. Although small RNAs isolated from aged mice have been revealed to activate TLR7 to induce senescence, the exact contributor from the fraction has not been disclosed^34^. Our findings revealed the first representative tRNA half molecule as a TLR7 agonist to induce cellular senescence. Similar to cGAS/STING triggers inflammaging, we demonstrated that the accumulated 5′-tRNA^Sec(NCA)^ half activates TLR7 to lead the release of inflammatory cytokines and promote cellular senescence, reinforcing the linkage between innate immune responses and inflammaging. Notably, mucosal TLR5 activation expands heathy lifespan in both male and female mice^53^. This differential impact of TLRs on aging underscores the complexity of the immune response in aging and highlights the need for targeted therapeutic strategies that modulate specific TLR pathways.

A series of aging hallmarks have been identified over the past three decades, which indeed serve as indicators reflecting the status and degree of aging to some extent^3^. However, they are mainly the consequences of the aging process rather than the root causes. Aging hallmark benefits various biological functions such as P16, whose accumulation promotes tissue-regeneration in young mice, indicating that it lacks of rationality to develop anti-aging therapeutics *via* targeting aging markers^54^. Therefore, targeting a direct driver of senescence, such as 5′-tRNA^Sec(NCA)^ half in this work, is more precise. Our results show that ASO-5′-tRNA^Sec(NCA)^ half effectively reduces senescence markers, increases telomere length, and extends the healthspan and lifespan of aged mice. Moreover, we revealed 5′-tRNA^Sec(NCA)^ half accumulation in humans during aging and confirmed the anti-aging effect of its ASO in human aged cells. This highlights the potential of applying ASO-5′-tRNA^Sec(NCA)^ half for aging intervention in clinics. Since oligonucleotide therapeutics including siRNA and ASO have exhibited huge potential in pharmacy market and clinical application, this broadens the application scenarios of small nucleic acid drugs, holding significant implications for the future development of the nucleic acid drug industry.

In addition to the blockage effect of ASO on 5′-tRNA^Sec(NCA)^ half, we also revealed that this ASO molecule could potentially function through RNAi pathway to suppress an aging-related gene, BCAT1, which was significantly downregulated in aged mice rather than adult mice in proteomics analysis. This conserved phenomenon has been verified in *C. elegans*, *D. rerio* and *M. musculus*, which might be an endogenous regulatory mechanism in response to aging during the process of biological evolution^43^. BCAT1 plays a vital role in promoting BCAAs (leucine, isoleucine and alanine) catabolism, whose high expressions lead to multiple pathological states such as gliomas, sepis, etc.^55,56^. Thus, BCAT1 inhibition blocks the catabolism of BCAAs to increase their expressions^57^. Indeed, BCAAs supplementation has been revealed to promote healthy aging such as extending the average lifespan of male mice, enhancing mitochondrial biogenesis in cardiac and skeletal muscles, and reducing oxidative damage^58^. However, increased BCAAs activate mTOR signaling pathway, in particular, leucine has been addressed as an agonist of mTOR^59^. High BCAAs dietary leads to an approximately 10% reduction in the lifespan of mice^60,61^. Notably, restriction of BCAAs dietary remarkably increases lifespan and healthspan in progeroid male mouse for 30%^62^. These findings suggest a risk when using BCAAs supplementation to intervene the aging process. Thus, it is essential to develop personalized regimens in line with the actual conditions of individual populations. Our findings provide the first evidence that BCAT1 inhibition exhibits the outcome of lifespan extension in both male and female mice, suggesting BCAT1 regulation might be a more promising and safer strategy to intervene aging. Future studies should pay more attention to the RNAi therapeutics targeting BCAT1 to investigate the anti-aging effect in mammals.

In summary, our study uncovers tRNA damage as a novel hallmark of aging, with 5′-tRNA^Sec(NCA)^ half playing a crucial role in inducing cellular senescence. ASO targeting 5′-tRNA^Sec(NCA)^ half not only effectively blocks the senescence-inducing effect of this tRNA half but also functions through the RNAi pathway to suppress BCAT1 mRNA, thereby extending the lifespan and healthspan of mice. However, several limitations exist. While our findings strongly suggest the anti-aging potential of targeting 5′-tRNA^Sec(NCA)^ half and BCAT1, further research is needed to explore whether other pathways are involved in the anti-aging effects of ASO. Additionally, given the complexity of aging, investigating the impact of this intervention on a broader range of aging-related parameters and in different mouse strains would provide more comprehensive insights. Future studies should also focus on translating these findings to human applications, evaluating the safety and efficacy of ASO-5′-tRNA^Sec(NCA)^ half in clinical settings. Overall, our work lays a solid foundation for further exploration of tRNA damage as a therapeutic target for aging and offers promising directions for the development of novel anti-aging strategies.

## Methods

### Chemicals and materials

All chemicals and materials used in this study were summarized in Supplementary Table S2.

### Animals

*D. melanogaster* was purchased from Qidong Fungene Biotechnology Co., Ltd. *C*. *elegans* was purchased from Shanghai Model Organisms Center. Wild-type 16-month-old C57BL/6J mice were purchased from Zhishan (Beijing) Institute of Health Medicine Co., Ltd. Wild-type 6- and 20-month-old Balb/c female mice were purchased from Biocytogen Jiangsu Co., Ltd. All animals were maintained in a controlled environment featuring a 12-hour light/dark cycle (lights on at 7:00) and provided with unlimited access to food and water. The Animal Ethical and Welfare Committee of Macau University of Science and Technology approved all experimental protocols under the ethical approval number MUSTARE-003-2020. All procedures for animal care and handling adhered to the National Institutes of Health Guidelines for the Care and Use of Laboratory Animals.

### tRNA mapping by UHPLC-MS

Total tRNAs (10 µg) were firstly digested by RNase T1 (50 U for 1 µg RNA) in 220 mM ammonium acetate buffer at 37C for 2 h. The mixtures were then incubated at 70C for 10 min to stop the reaction. After centrifugation (10,000×g for 1 min), the supernatants contained hydrolysates were collected for UHPLC-MS/MS analysis. For tRNA mapping, tRNA MODOMICS database (https://genesilico.pl/modoomics/sequences/) was used to establish a mouse tRNA RNase T1 fragments database. The molecular formula of tRNA fragments was calculated using a simple program written by R studio software. The identification of tRNA fragments was performed using Agilent MassHunter Workstation for automatic matching of tRNA spectra, with a mass accuracy of less than 5 ppm between the theoretical and calculated masses of multi-charged ions. Count the peak area of each identified tRNA fragment.

### Proteomic analysis

Samples were homogenized by MP FastPrep-24 homogenizer, and mixed with SDT buffer (4% SDS, 100 mM Tris-HCl, pH=7.6). The lysates were further sonicated and boiled for 15 min, followed by centrifuged at 14,000 ×g for 40 min, the supernatant was quantified with the BCA Protein Assay Kit (Roche). Proteins were separated on 10% SDS-PAGE gel. Protein bands were visualized by Coomassie Blue R-250 staining. DTT (with the final concentration of 40 mM) was added to each sample respectively and mixed at 600 rpm for 1.5 h (37C). After the samples cooled down to room temperature, IAA was added with the final concentration of 20 mM into the mixture to block reduced cysteine residues and the samples were incubated for 30 min in darkness, followed by transferred to the filters (Microcon units, 10 KDa), respectively. The filters were washed with 100 μL of UA buffer and with 100 μL of NH_4_HCO_3_ buffer (25 mM). Finally, trypsin was added to the samples (the trypsin: protein (wt/wt) ratio was 1:50) and incubated at 37C for 15-18 h (overnight), and the resulting peptides were collected as a filtrate. The peptides of each sample were desalted on C_18_ Cartridges (Empore™ SPE Cartridges MCX, 30 UM, Waters), concentrated by vacuum centrifugation and reconstituted in 40 µL of 0.1% (v/v) formic acid. The peptide content was estimated by UV light spectral density at 280 nm. For DIA experiments, iRT (indexed retention time) calibration peptides were spiked into the sample.

The peptides from each sample were analyzed by Orbitrap^TM^ Astral^TM^ mass spectrometer (Thermo) connected to an Vanquish Neo system liquid chromatography (Thermo) in the data-independent acquisition (DIA) mode. Precursor ions were scanned at a mass range of 380-980 m/z, MS^1^ resolution was 240,000 at 200 m/z, Normalized AGC Target: 500%, Maximum IT: 5 ms. 299 windows were set for DIA mode in MS^2^ scanning, Isolation Window: 2 m/z, HCD Collision Energy: 25 ev, Normalized AGC Target: 500%, Maximum IT: 3 ms. DIA data were analyzed with DIA-NN 1.8.1. Main software parameters were set as follows: enzyme is trypsin, max missed cleavages is 1, fixed modification is carbamidomethyl (C), dynamic modification is oxidation (M) and acetyl (Protein N-term). All reported data were based on 99% confidence for protein identification as determined by false discovery rate (FDR) ≤ 1%.

### tRNA half enrichment

Samples were mixed with guanidinium isothiocyanate buffer (4 M) for homogenization and lysis on ice for 20 min. After centrifugation at 10,000 ×g for 10 min, supernatant was collected and mixed with poly-arginine modified hydroxyapatite, followed by incubation for 30 min on ice. Subsequently, the mixture was centrifuged and the precipitates were washed with RNase-free water followed by centrifugation to remove unspecific binding chemicals. The precipitates were mixed with 0.4 M of NaH_2_PO_4_ buffer (pH=7.5) to release small RNAs, which were further desalted by ultrafiltration column (3 KDa).

Ion-pair chromatographic method was employed to separate tRNA halves from small RNAs. Separation was carried out on an ACQUITY UPLC OST C_18_ Column (2.1×100 mm i.d., 1.7 μm, Waters) at 60C. The flow rate was set at 0.2 mL/min and sample injection volume was 20 μL. Gradient elution with (A) 100 mM hexafluoro-2-propanol (HFIP)+15 mM trimethylamine (TEA) and (B) 50% MeOH in A was 0-1.5 min, 2%B, 1.5-8.3 min, 2-28%B, 8.3-16.5 min, 28-34%B, followed by washing with 80% B and equilibration with 2%B. The fraction of tRNA half was obtained and freeze-dried using a Speed-Vac system RVC 2-18 (Marin Christ, Germany).

### Clinical samples collection

White blood cells (WBCs) were collected from a cohort of 69 healthy volunteers aged 20 to 70 years. The study was conducted with the assistance of Guangzhou KingMed Diagnostics Group Co., Ltd. Ethical approval for the study was obtained, and informed consent was acquired from all participants prior to sample collection. Peripheral blood samples were drawn from each volunteer using standard venipuncture techniques. Approximately 5 mL of blood was collected into EDTA-coated tubes to prevent coagulation. Following collection, the blood samples were immediately processed to isolate WBCs. WBCs were isolated using a Ficoll-Paque density gradient centrifugation method. Briefly, blood samples were diluted with an equal volume of phosphate-buffered saline (PBS) and carefully layered over Ficoll-Paque PLUS (GE Healthcare). The samples were then centrifuged at 400 ×g for 30 min at room temperature. The WBC layer was carefully aspirated and washed twice with PBS to remove any remaining plasma and platelets. The purified WBCs were then resuspended in PBS and counted using a hemocytometer. The isolated WBCs were aliquoted and stored at -80°C until further isolated for tRNA half analysis.

### Cell culture

RAW264.7 mouse macrophage cell line and TCMK-1 mouse kidney tubule epithelial cell line purchased from American Type Culture Collection was cultured in Dulbecco’s Modified Eagle Medium (DMEM) supplemented with 10% fetal bovine serum (FBS), 100 U/mL penicillin, and 100 µg/mL streptomycin. 2BS human fetal lung fibroblasts cell line purchased from American Type Culture Collection was cultured in Minimum Essential Medium (MEM) supplemented with 10% fetal bovine serum (FBS), 1% non-essential amino acids (NEAA), 100 U/mL penicillin, and 100 µg/mL streptomycin. All cell lines were maintained at 37°C in a humidified atmosphere containing 5% CO_2_.

### Cellular senescence model

For chemo-inducing aged cells, TCMK-1 cells were seeded and allowed to grow overnight. Subsequently, medium were replaced with culture medium containing D-galactose at a final concentration of 20 mg/mL. The cells were cultured for another 72 h to set a cellular senescence model. For naturally aged cells, 2BS cells were normally cultured to 37^th^ passage to set a cellular senescence model.

### Nitric oxide release determination

In brief, RAW264.7 cells were seeded at a density of 1×10^5^ cells per well in 96-well microplates and incubated overnight. The cells were then treated with t-halves fraction transfected by Lipofectamine RNAiMAX reagent, as well as lipopolysaccharide (LPS) used as a positive control. After incubation for 24 h, an equal volume of culture medium served as the blank control. After the treatment period, 75 μL of the supernatant from each well was transferred to a new 96-well plate. An equal volume of Griess reagent was added to each well, and the reaction was allowed to proceed at room temperature for 15 min. The absorbance was then measured at 540 nm using a microplate reader. NO production was determined by calculating the ratio of absorbance values between the treatment groups and the LPS-treated group.

### Purification and characterization of 5**′**-tRNA^Sec(NCA)^ half

5′-tRNA^Sec(NCA)^ half purification was carried out according to our previous method^34^. In brief, tRNA half fraction obtained from kidneys of aged were firstly separated by LC-MS-aided ion-pair chromatography to afford the fraction containing the target oligonucleotide of 5′-tRNA^Sec(NCA)^ half. After freeze-dried, the samples were dissolved by RNase-free water and injected to weak anion-exchange chromatography to afford 20 fractions, which were then assayed by LC-MS analysis. The fraction contained 5′-tRNA^Sec(NCA)^ half was further digested by RNase T1 and characterized by UHPLC-MS/MS.

For oligonucleotides analysis, an Agilent UHPLC 1290 system (Agilent Technologies, Santa Clara, CA, U.S.A.) was applied for UHPLC-MS/MS analysis of RNase T1 digestion products, which was equipped with a vacuum degasser, a quaternary pump, an autosampler, a diode array detector, and an Agilent ultrahigh definition 6545 Q-TOF mass spectrometer. An ACQUITY UPLC OST C_18_ column (2.1×100 mm i.d., 1.7 μm, Waters, Massachusetts, U.S.A.) was maintained at 60C for chromatographic separation. The flow rate was set at 0.2 mL/min and sample injection volume was 20 μL. Gradient elution with (A) 100 mM 1,1,1,3,3,3-hexafluoto-2-propanol (HFIP)+15 mM trimethylamine (TEA) and (B) 50% MeOH in A was 0-1.5 min, 2%B, 1.5–8.3 min, 2%–28%B, 8.3–16.5 min, 28%–34%B, followed by washing with 80%B and equilibrating with 2%B. ESI conditions were set as follows: gas temperature 320C, spray voltage 3.5 kV, sheath gas flow and temperature were set as 12 L/min and 350C, respectively. For MS experiment, samples were analyzed in negative mode over an m/z range of 500 to 3200. For MS/MS experiment, samples were analyzed in negative mode over an m/z range from 300 to 2000. The information of sequence and modification were obtained from the collision induced dissociation analysis of target oligonucleotides.

### CCK-8 assay

Briefly, 1×10^3^ cells per well in 96-well microplates were seeded and incubated overnight. After 20 h, RNA samples (purchased from Biosyntech, China. Supplementary Table S3) encapsulated by liposomes were added. At the day 1, 2, 3, 4, 5, and 6, 10 μL of CCK8 solution was added to each well containing 100 μL of culture medium. The plates were then incubated at 37C in a humidified atmosphere with 5% CO_2_ for 4 h. The absorbance of each well was measured at 450 nm using a microplate reader. Cell viability was calculated as a percentage of the absorbance of treated cells relative to the control cells.

### SA-**β**-gal staining

2×10^5^ cells of TCMK-1 or 2BS per well were seeded in 6-well microplates. After RNA samples treatment, SA-β-gal activity was assayed by Senescence β-Galactosidase Staining Kit according to the manufactures’ protocol (Beyotime, China).

### ROS determination

The intracellular ROS level was measured using the ROS assay kit (Beyotime, China). Briefly, TCMK-1 or 2BS was seeded in 6-well plates at a density of 5×10^4^ cells/well. After drug treatment, DCFH-DA was diluted with serum-free culture medium at a ratio of 1:1000 and incubated at 37C for 20 min. After incubation, the cells were washed 3 times with serum-free cell culture medium for 3 min each time. The entire experimental process was completed in the dark. Representative images were acquired using an inverted fluorescence microscope (Leica Olympus ix73, Germany) and fluorescence normalization was performed.

### RNA isolation and RT-qPCR

Total RNA extraction was carried out according to the manufactures’ guideline of TRIzol reagent. The RNA was then subjected to reverse transcription using the GoScript Reverse Transcription System (Promega, U.S.A.). Quantitative real-time PCR was performed on a ViiATM 7 system (Life Technologies, U.S.A.) using the GoTaq® qPCR Master Mix (Promega, U.S.A.) according to the manufacturers’ instructions. The PCR primers were commercially obtained from BGI (China, Supplementary Table S4). The data are presented as the average fold change values of duplicates and normalized to GAPDH expression using the 2−ΔΔCT method. Each experiment was conducted in triplicate, and the results are reported as means ± standard deviation (SD).

### Telomere length measurement

Genomic DNA was extracted using the Universal Genomic DNA Purification Mini Spin Kit (Beyotime) according to the manufacturer’s instructions. DNA concentration was measured with a NanoDrop 2000 spectrophotometer (ThermoFisher Scientific, Waltham, MA, USA). Telomere length (TL) was assessed using real-time quantitative PCR^63,64^. The relative TL was determined by comparing the ratio of T repeat copy number to S copy number, expressed as the telomere length (T/S) ratio.

### Transcriptomic sequencing

Total cellular RNA was extracted using Trizol Reagent (Thermo) according to the manufacturer’s instructions. RNA quality and quantity were evaluated using the K5500 spectrophotometer (BeijingKaiao, China) and the Agilent 2200 TapeStation (Agilent Technologies). mRNA enrichment was performed using oligodT as per the NEBNext® Poly(A) mRNA Magnetic Isolation Module (NEB, USA). The enriched mRNA was fragmented to approximately 200 bp. First and second strand cDNA synthesis, adaptor ligation, and low-cycle enrichment were conducted using the NEBNext® Ultra™ RNA Library Prep Kit for Illumina. The purified library products were assessed with the Agilent 2200 TapeStation and quantified using Qubit (Thermo). Paired-end sequencing with 150 bp reads was carried out on an Illumina platform (Illumina, USA).

### ELISA assay of interferon

ELISA assay was carried out using Mouse and Human IFN-β ELISA Kit (Multi Sciences, China) according to manufactures’ instructions.

### Western blotting

Cells or tissues were lysed and proteins were extracted using RIPA buffer (Cell Signaling Technology, U.S.A.) that was supplemented with protease and phosphatase inhibitor cocktails (Roche, Switzerland). Protein concentrations were determined using a BCA protein assay (Thermo). Denatured proteins were separated using 10% SDS-PAGE and then transferred onto nitrocellulose membranes. These membranes were subsequently blocked with 5% bovine serum albumin (Thermo) for 2 h. After blocking, the membranes were incubated overnight with primary antibodies against. Following a wash with TBST, the membranes were incubated with a fluorescent secondary antibody for detection purposes. Band intensities were quantified using ImageJ software and normalized to the intensities of GAPDH. Each experiment was conducted in triplicate, and the results are presented as means ± standard deviation (SD).

### RNA pull-down

Harvested TCMK-1 cells were washed twice with ice-cold PBS, add cell lysis buffer with protease and phosphatase inhibitors, incubate on ice for 15-30 min, and centrifuge at 12,000 ×g for 15 min at 4C to obtain the cell lysate supernatant. Wash streptavidin magnetic beads three times with binding buffer, add the cell lysate for 1-2 hours of incubation at 4C with gentle rotation to pre-clear, then transfer the supernatant. Add the biotinylated RNA (5′-tRNA^Sec(NCA)^ half, scrambled 5′-tRNA^Sec(NCA)^ half and ASO-5′-tRNA^Sec(NCA)^ half) to the pre-cleared lysate and incubate at room temperature for 1-2 h. Subsequently, add streptavidin magnetic beads and incubate at 4C for 1-2 h. Wash the beads three to five times with washing buffer. Elute the bound proteins with an elution buffer at room temperature for 10-15 min, separate the eluate, and analyze the proteins by techniques like western blotting or mass spectrometry.

### Paraquat-model to determine anti-aging effect

6 to 8-week-old Balb/c female mice were intraperitoneally (i.p.) injected with paraquat (Sigma) at the dose of 50 mg/kg at day 1. LNP encapsulated ASO-5′-tRNA^Sec(NCA)^ half (2.5 and 1.25 mg/kg) or PBS were intravenously injected (i.v.) to the mice at 2 h and 10 h post the paraquat modeling. Animals were sacrificed at Day 3 to harvest their tissues.

### Lifespan studies for natural aging

A total of 40 mice were randomly divided into two groups: the LNP group (n=20) and the ASO-5′-tRNA^Sec(NCA)^ half group (2.5 mg/kg, n=20). Starting at 16-month-old age, all mice received intravenous injections of either LNP or LNP encapsulated ASO-5′-tRNA^Sec(NCA)^ half every two weeks until death. At 20-month-old age, six mice from each group were randomly selected to collect blood through intravenous for DNA methylation clock determination. At 23-month-old age, eight mice from each group were randomly selected to test their behavior. At 24-month-old age, eight mice from each group were randomly selected to test their bone mass and muscle. At 26-month-old age, five mice from each group were randomly selected to analyze biochemical markers. The remaining mice were allowed to die naturally, and the time of death was recorded to calculate lifespan.

### Determination of aging markers in WBCs of natural mice

Blood collection from the fundus of the eye was performed to obtain 100 μL of whole blood from naturally aged mice. This method involved gently restraining the mouse and using a microcapillary tube to carefully collect a small volume of blood from the punctured fundus. Subsequently, total mRNA were extracted from whole blood by using Magnetic Tissue/Cell/Blood Total RNA Kit (TIANGEN Biotech, China) according to the manufactures’ protocol, followed by RT-qPCR analysis.

### Reduced representation bisulfite sequencing

Genomic DNA was extracted and subjected to RRBS library preparation using the Acegen Rapid RRBS Library Prep Kit (Acegen, Cat. No. AG0422) according to the manufacturer’s protocol. In brief, 100 ng of genomic DNA was digested with MspI, end-repaired, 3’-dA-tailed and ligated to 5-methylcytosine-modified adapters. After bisulfite treatment, the DNA was amplified with 12 cycles of PCR using Illumina 8-bp dual index primers. Size selection was performed to obtain DNA fractions of MspI-digested products in the range of 100-350 bp using a dual-SPRI® protocol according to the manufacturer’s protocol. The constructed RRBS libraries were then analyzed by Agilent 2100 Bioanalyzer and finally sequenced on Illumina platforms using a 150×2 paired-end sequencing protocol.

### Calculation of DNA methylation age

Initially, the result files of the GSE80672 dataset were downloaded from the GEO database^65^. Subsequently, 163 appropriate samples, including those from specific datasets and wild-type mice, were carefully selected. From these samples, the methylation levels and coverage depths of 90 sites were extracted. Data filtering was then performed based on a coverage depth greater than 4X, and missing values were interpolated using KNN. The dataset was divided into a training set and a validation set in an 8:2 ratio to construct an elastic network model. After model evaluation and analysis of the correlation between predicted and actual ages, the predicted ages were fitted and corrected against the actual ages. Finally, the same coverage criteria were applied to the tested samples for filtering and interpolation, and the prediction model was used for prediction and correction to obtain the predicted age results of each sample using the calculation formula below, providing a solid foundation for further research.

Age^met^=a*Age^b^+c

*Age^met^*stands for the predicted DNA methylation age of tested mice

*Age* stands for actual age of tested mice a=2.9727, b=0.6818, c=-2.2335

### Animal behavior test

#### Open field test

The open field test was conducted to assess general locomotor activity and anxiety-like behavior. Mice were placed individually in the center of a square open field arena (50 cm×50 cm) with walls 40 cm high. The arena was divided into equal quadrants by lines drawn on the floor. Each mouse was allowed to explore the arena for 10 min, and their movements were recorded using a video tracking system. Parameters such as total distance traveled, time spent in the center versus the periphery, and the number of entries into the center were analyzed to evaluate locomotion and anxiety levels. The arena was cleaned with 70% ethanol between trials to eliminate olfactory cues.

### Y-maze test

The Y-maze test was performed to evaluate spatial working memory and exploratory behavior. The apparatus consisted of three arms (each 35 cm long, 5 cm wide, and 15 cm high) arranged in a Y-shape. Mice were placed at the end of one arm and allowed to explore the maze freely for 8 minutes. The sequence and number of arm entries were recorded. An entry was defined as all four paws within an arm. The percentage of spontaneous alternations (consecutive entries into all three arms without repetition) was calculated to assess working memory. The maze was cleaned with 70% ethanol between trials to remove scent traces.

### Grip strength test

The grip strength test was used to measure forelimb muscle strength. Mice were gently held by the tail and allowed to grasp a horizontal metal bar connected to a grip strength meter (Bioseb, France). Once the mouse had a firm grip, it was gently pulled backward until it released the bar. The peak force exerted by the forelimbs was recorded in grams. Each mouse performed five trials, and the average grip strength was calculated. The apparatus was cleaned with 70% ethanol between trials to ensure consistency.

### Rotarod test

The rotarod test was conducted to assess motor coordination and balance. Mice were placed on a rotating rod (rotarod apparatus, Ugo Basile) with an initial speed of 4 rpm, which gradually increased to 40 rpm over 5 minutes. The latency to fall off the rod was recorded for each mouse. Mice underwent three trials per day for three consecutive days, with at least 15 min of rest between trials. The average latency to fall was calculated for each mouse. The apparatus was cleaned with 70% ethanol between trials to maintain hygiene.

### Tail suspension test

The tail suspension test was performed to evaluate depressive-like behavior. Mice were suspended by the tail using adhesive tape attached to a horizontal bar, ensuring that their heads were approximately 20 cm above the surface. Each mouse was suspended for 6 min, and the duration of immobility was recorded. Immobility was defined as the absence of any limb or body movements except those necessary for breathing. The test was conducted in a quiet room with minimal disturbances. The apparatus was cleaned with 70% ethanol between trials to eliminate any residual odors.

### Dual-energy X-ray absorptiometry

Dual-energy X-ray absorptiometry (DXA) scans were carried out by employing a KUBTEC PARAMETER 3D X-ray cabinet in conjunction with DIGIMUS software (KUB Technologies, Stratford, CT, USA). These scans were obtained when the mice were under isoflurane anesthesia. For the quantitative bone analyses, the mid-diaphysis of both the left and right tibia was selected. This region is positioned exactly halfway between the tibial plateau and the tibiofibular junction. The bone mineral density (BMD, measured as g hydroxyapatite/cm²) and bone mineral content (BMC, measured as g hydroxyapatite) were determined using the software provided by the manufacturer.

### Histological investigations

Mouse tissues were fixed in 4% paraformaldehyde (PFA) and subsequently embedded in paraffin. For immunohistochemistry (IHC), paraffin-embedded sections were deparaffinized in xylene and rehydrated through a graded ethanol series. Antigen retrieval was performed by incubating the sections in 10 mM sodium citrate buffer (pH 6.0) at 95C for 30 min. The sections were then treated with 3% hydrogen peroxide for 15 min to quench endogenous peroxidase activity, followed by three washes in phosphate-buffered saline (PBS), each lasting 3 min. Non-specific binding sites were blocked with 5% bovine serum albumin (BSA). The sections were incubated overnight at 4°C with primary antibodies against TLR7, P16 and P21. After thorough washing with PBS, the sections were incubated with a GTVison secondary antibody. Detection of antibody complexes was achieved using 3,3’-diaminobenzidine (DAB) as the chromogen, followed by counterstaining with Mayer’s Hematoxylin. To evaluate potential side effects of ASO-5′-tRNA^Sec(NCA)^ treatment, 3-5 μm paraffin sections were also subjected to hematoxylin and eosin (H&E) staining.

### Bioinformatics analysis

Target prediction was conducted using miRanda and TargetScan according to our previous report^66^. miRanda utilizes RNA secondary structure and free energy to identify any seed type, while TargetScan predicts biologically significant sites based on mRNA-miRNA expression profiles and conservation scores, specifically targeting m8, 7mer-m8, and 7mer-a1 seed types. Combining these algorithms leveraged their strengths for improved prediction and visualization. The 2D structure of binding sites, including position, type, and base pairing, was detailed, with critical pairings highlighted. Local AU content and site position within the UTR were also considered for accessibility and favorability. Filtering parameters included structure score and free energy for miRanda, and context+score for TargetScan, ensuring robust and accurate miRNA target site predictions.

### Dual-luciferase reporter assay

The pmiR-RB-REPORT vector (Ribobio, China) was utilized to introduce the wild-type or mutant ASO-5′-tRNA^Sec(NCA)^ half binding sites into the Xhol and Notl restriction sites within the 3’ UTR of the BCAT1 sequence. HEK293T cells were co-transfected with the luciferase reporter vectors and either the ASO-5′-tRNA^Sec(NCA)^ half or its scrambled sequence using Lipofectamine RNAiMAX Transfection Reagents (Thermo). After transfection for 48 h, the Firefly and Renilla luciferase activities were quantified using the Dual-Glo Luciferase Assay System (Promega) and the GLOMAX 96 spectrophotometer (Promega), following the provided instructions. Each experiment was repeated three times, and the results are presented as means ± standard deviation (SD).

### TCGA data analysis

RNA sequencing data of clinical samples for various cancers were retrieved from TCGA database (https://portal.gdc.com). The latest release of GTEx database was utilized through GTEx data portal website (https://www.gtexportal.org/home/datasets). R software v4.0.3 (R Foundation for Statistical Computing, Austria) was employed for statistical analysis. *P* value <0.05 is considered as statistical significance.

## Acknowledgments

We thank Dr. Tao Chen from Guangzhou KingMed Diagnostics Group Co., Ltd. for his assistance to collect clinical blood samples. This work was financially funded by The Science and Technology Development Fund, Macau SAR (File no. 006/2023/SKL and 0001/2023/AKP to Z.-H. Jiang). This work was partially supported by Ruina (Zhuhai Hengqin) Biotechnology.

## Author contributions

Z.-H.J. and K.-Y.C. conceived the study. K.-Y.C., L.-B.B., D.Z., Y.-H.Z., R.-Z.G. T.-M.Y, Y.P. and Y.-T.C. performed the experiment. L.-B.B. and D.Z. contributed to the materials. K.-Y.C., L.-B.B., D.Z. and Y.-H.Z. analyzed the data. K.-Y.C. and Z.-H.J. contributed to the discussion and wrote the manuscript.

