## Supplemental Materials for "Targeted intervention of senescence induced by a 5′ half fragment of tRNA^Seca(NCA)^ with antisense oligonucleotide extends healthspan and lifespan in mice"

This file includes:

1. Supplementary Tables S1-S4
2. Supplementary Figures S1-S8

**Table S1. MS$^2$ data of RNase T1 digestion products of 5′-tRNA$^{Sec(NCA)}$ half purified from kidney of aged mice.**

| Product | RNase T1 digestion products' sequence (5'-3') | Molecular formula | Measured m/z for molecular ion | *m/z* | Calculated m/z for molecular ion | Difference[a] |
|---|---|---|---|---|---|---|
| 1 | CCCGp | $C_{37}H_{50}N_{14}O_{29}P_4$ | 638.0899 | $[M-2H]^{2-}$ | 638.0831 | 0.0068 |
| 2 | AUGp | $C_{29}H_{37}N_{12}O_{22}P_3$ | 997.1357 | $[M-H]^{-}$ | 997.1280 | 0.0077 |
| 3 | AUCCUCAGp | $C_{75}H_{96}N_{28}O_{57}P_8$ | 1273.1629 | $[M-2H]^{2-}$ | 1273.1610 | 0.0019 |
| 4 | UGp | $C_{19}H_{25}N_7O_{16}P_2$ | 668.0773 | $[M-H]^{-}$ | 668.0755 | 0.0018 |
| 5 | UCUGp | $C_{37}H_{48}N_{12}O_{31}P_4$ | 639.0735 | $[M-2H]^{2-}$ | 639.0671 | 0.0064 |
| 6 | CAGp | $C_{29}H_{38}N_{13}O_{21}P_3$ | 996.1468 | $[M-H]^{-}$ | 996.1440 | 0.0028 |
| 7 | CU | $C_{18}H_{24}N_5O_{13}P_1$ | 548.1055 | $[M-H]^{-}$ | 548.1030 | 0.0025 |

[a] Difference = (Measured mass)-(Calculated mass)

**Table S2. Reagents and resources.**

| Reagents or Resources | Source | Identifier |
|---|---|---|
| **Antibodies** | | |
| Mouse monoclonal anti-TLR7 | Novus Biologicals | Cat# NBP2-27332 |
| Rat monoclonal anti-P16 | Abcam | Cat# ab241543 |
| Rabbit polyclonal anti-P21 | ProteinTech | Cat# 28248-1-AP |
| Rabbit polyclonal anti-IL6 | ProteinTech | Cat# 26404-1-AP |
| Rabbit polyclonal anti-BCAT1 | Abcepta | Cat# AP10147c |
| Rabbit monoclonal anti-GAPDH | Cell Signaling Technology | Cat# 5174 |
| IRDye 800CW Goat anti-Rabbit IgG | LI-COR Biosciences | Cat# 926-32211 |
| **Chemicals** | | |
| TRIzol | Thermo Fisher | Cat# 15596026 |
| RNase T1 | Thermo Fisher | Cat# EN0542 |
| BeyoFast™ SYBR Green qPCR Mix | Beyotime | Cat# D7260 |
| Cell Counting Kit-8 | Beyotime | Cat# C0037 |
| LPS | Beyotime | Cat# S1732 |
| RIPA | Cell Signaling Technology | Cat# #9806 |
| Low range ssRNA ladder | New England Biolabs | Cat# N0364S |
| microRNA ladder | New England Biolabs | Cat# N2102S |
| LNP | Scindy Pharmaceutical | Cat# SDR8002 |
| D-galactose | Macklin | Cat# D796635 |
| Paraquat | Macklin | Cat# M919425 |
| Griess reagent | Sigma | Cat# G4410 |
| Commercial kits | | |
| RBC Lysis Buffer (10X) | BioLegend | Cat# 420302 |
| RevertAid RT Reverse Transcription Kit | Thermo Fisher | Cat# K1691 |
| Pierce BCA Protein Assay Kits | Thermo Fisher | Cat# A65453 |
| Pierce Magnetic RNA-Protein Pull-Down Kit | Thermo Fisher | Cat# 20164 |
| Streptavidin-coated magnetic beads | Beaver Biotechnology | Cat# 22308-1 |
| mirVana miRNA Isolation Kit | Thermo Fisher | Cat# AM1561 |
| Senescence β-Galactosidase Staining Kit | Beyotime | Cat# C0602 |
| Universal Genomic DNA Purification Mini Spin Kit | Beyotime | Cat# D0063 |
| Reactive Oxygen Species Assay Kit | Beyotime | Cat# S0033S |
| Mouse IFN-β ELISA Kit | Multi Sciences | Cat# EK2236 |
| Human IFN-β ELISA Kit | Multi Sciences | Cat# EK1236 |
| Mouse IFN-gamma ELISA kit | Multi Sciences | Cat# EK280 |
| Human IFN-gamma ELISA kit | Multi Sciences | Cat# EK180 |
| Mouse IL-6 ELISA Kit | Multi Sciences | Cat# EK206 |
| Mouse telomerase,TE ELISA Kit | CUSABIO | Cat# CSB-E08022m |
| Triglyceride assay kit | Nanjing Jiancheng Bioengineering Institute | Cat# A110-1-1 |
| Total cholesterol assay kit | Nanjing Jiancheng Bioengineering | Cat# A111-1-1 |

| | | |
|---|---|---|
| | Institute | |
| High-density lipoprotein cholesterol assay kit | Nanjing Jiancheng Bioengineering Institute | Cat# A112-1-1 |
| Low-density lipoprotein cholesterol assay kit | Nanjing Jiancheng Bioengineering Institute | Cat# A113-1-1 |
| Creatinine (Cr) Assay kit | Nanjing Jiancheng Bioengineering Institute | Cat# C011-2-1 |
| Glucose kit | Nanjing Jiancheng Bioengineering Institute | Cat# A154-1-1 |
| Estradiol 2 Assay Kit | Nanjing Jiancheng Bioengineering Institute | Cat# H102-1-2 |
| Follicle stimulating hormone Assay Kit | Nanjing Jiancheng Bioengineering Institute | Cat# H101-1-2 |
| Progesterone Assay Kit | Nanjing Jiancheng Bioengineering Institute | Cat# H089-1-2 |
| Anti-Mullerian Hormone assay kit | Nanjing Jiancheng Bioengineering Institute | Cat# H324-1-2 |

### Table S3. Oligonucleotides used in this study.

| Gene name | Forward (5'-3') | Reverse (5'-3') |
|---|---|---|
| 5′-tRNA$^{Sec(NCA)}$ half | GCCCGGAUGAUCCUCAGUGGUC UGGGGUGCAGGCU | / |
| Scrambled 5′-tRNA$^{Sec(NCA)}$ half | AGCCGUACCCCCGAACCCAGUGA AGCUUAGGGCCC | / |
| ASO-5′-tRNA$^{Sec(NCA)}$ half | AGCCUGCACCCCAGACCACUGAG GAUCAUCCGGGC | / |
| siTLR7 for mice | GGAGCACACAAAGGUCAAATT | UUUGACCUUUGUGUGCUC CTT |
| siTLR7 for human | GCCUUGAGGCCAACAACAUUU | AAAUGUUGUUGGCCUCAA GGC |

### Table S4. Primers for quantitative real-time PCR.

| Gene name | Forward (5'-3') | Reverse (5'-3') |
|---|---|---|
| For mice | | |
| P16 | CGCAGGTTCTTGGTCACTGT | TGTTCACGAAAGCCAGAGCG |
| P21 | CCTGGTGATGTCCGACCTG | CCATGAGCGCATCGCAATC |
| IL6 | TAGTCCTTCCTACCCCAATTTCC | TTGGTCCTTAGCCACTCCTTC |
| CCL2 | TTAAAAACCTGGATCGGAACCAA | GCATTAGCTTCAGATTTACGGGT |
| TLR7 | CCACATTCACTCTCTTCATTGG | GGTCAAGAACTTCCAGCCTG |
| MYD88 | ATACGCAACCAGCAGAAACAG | TATCATTGGGGCAGTAGCAGA |
| RIG-I | AAGAGCCAGAGTGTCAGAATCT | AGCTCCAGTTGGTAATTTCTTGG |
| MDA5 | GCCTGGAACGTAGACGACAT | TGGTTGGGCCACTTCCATTT |
| BCAT1 | GAAGGAGATGTTCGGCTCAG | TCAGTCAGCTTTCCCAGGAT |
| DICER | GGTCCTTTCTTTGGACTGCCA | GCGATGAACGTCTTCCCTGA |
| ANG | GCAACCAGCGCCGAAAT | GGCACATTGCCCATGTTGA |
| RNase P | GCTCTACGCTTGGGCAGAC | TGAATTGGGTTATGAGGCCCG |
| GPX1 | GGTTCGAGCCCAATTTTACA | CCCACCAGGAACTTCTCAAA |
| TXNRD1 | AGTCACATCGGCTCGCTGAACT | GATGAGGAACCGCTCTGCTGAA |
| SelH | CCTTATTCCACCAACGCGCCA | GCGTCAGCTCGTACAATGCTC |
| SelO | TGACACTGAGTTCCAAAGGCAC | GTTAGTGAAGTCAGCACCAGTCAG |
| TELB | CGGTTTGTTTGGGTTTGGGTTTGGGT TTGGGTTTGGGTT | GGCTTGCCTTACCCTTACCCTTACCCTT ACCCTTACCCT |
| 36B4 | ACTGGTCTAGGACCCGAGAAG | TCAATGGTGCCTCTGGAGATT |
| GAPDH | TGAAGGTCGGTGTGAACGGATTTGG | ACGACATACTCAGCACCGGCCTCAC |
| For human | | |
| P16 | CCGTGGACCTGGCTGAGGAG | CGGGGATGTCTGAGGGACCTTC |
| P21 | CGATGGAACTTCGACTTTGTCA | GCACAAGGGTACAAGACAGTG |
| IL6 | CACTGGTCTTTTGGAGTTTGAG | GGACTTTTGTACTCATCTGCAC |
| CCL2 | GTGTCCCAAAGAAGCTGTGATCT | TGTCCAGGTGGTCCATGGA |
| TLR7 | CACAGCCGTCCCTACTGTTT | CTGTCAGCGCATCAAAAGCA |
| MYD88 | CGGATGGTGGTGGTTGTCTC | CCTCAGGATGCTGGGGAACT |
| RIG-I | AGTGAGCATGCACGAATGAA | GGGATCCCTGGAAACACTTT |
| MDA5 | TGTATTCATTATGCTACAGAACTG | ACTGAGACTGGTACTTTGGATTCT |
| BCAT1 | TACCGGAGAAGGAGGATCAA | TGGTGTGACTATTAGGTCTTTAGCC |
| TELB | CGGTTTGTTTGGGTTTGGGTTTGGGT TTGGGTTTGGGTT | GGCTTGCCTTACCCTTACCCTTACCCTT ACCCTTACCCT |
| 36B4 | CAGCAAGTGGGAAGGTGTAATCC | CCCATTCTATCATCAACGGGTACAA |
| GAPDH | TGAAGGTCGGAGTCAACGGATT | CGTTCTCAGCCTTGACGGT |

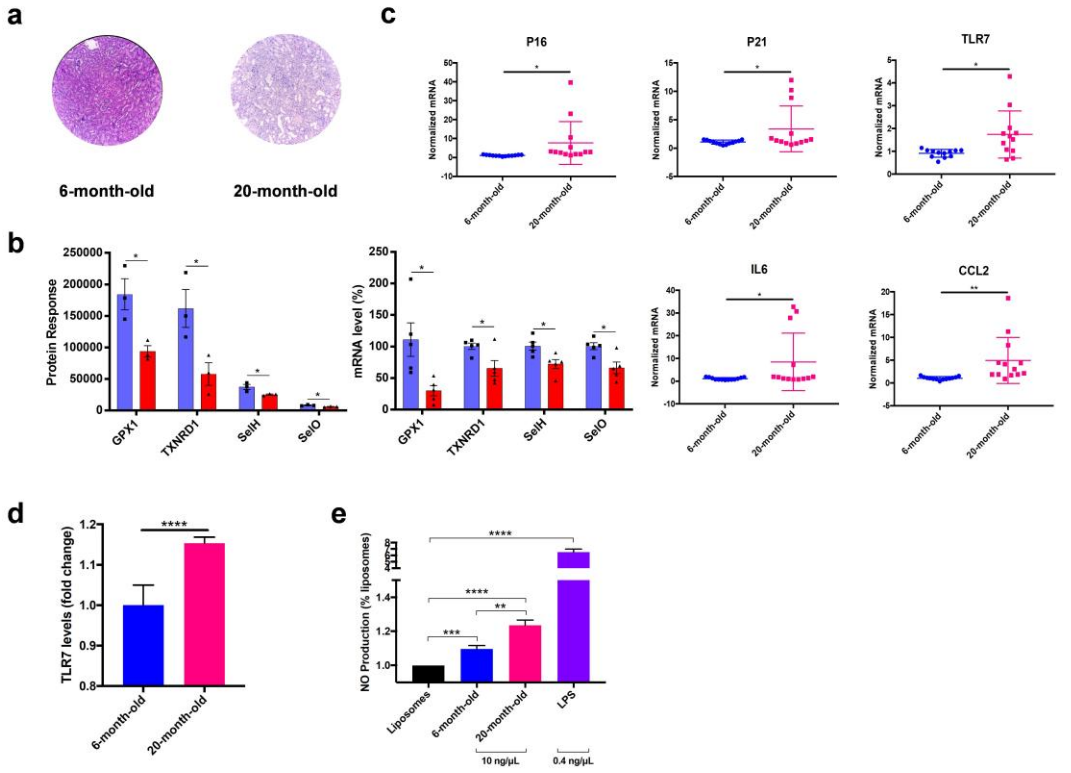


**Figure S1. Aged mice (20-month-old) exhibited significant high level of senescence than adult mice (6-month-old mice)**. **a**, H&E staining of kidneys from adult and aged mice. **b**, Expressions of GPX1, TXNRD1, SelH, and SelO in adult (Blue) and aged (Red) mice in mRNA and protein level. **c**, mRNA expressions of aging hallmarks (P16, P21), inflammation cytokines (IL6) and RNA sensor (TLR7) in kidneys from adult and aged mice. **d**, t-half fraction isolated from aged mice significantly increased TLR7 expressions and (**e**) promoted the release of nitric oxidation in RAW264.7 cells rather than that from adult mice. Data presented in **b**, **c** are presented as mean ± S.EM. Data presented in **d**, **e** are presented as mean ± S.D. Statistical test: two-tailed unpaired t-test. *$P$<0.05; **$P$<0.01; ***$P$<0.001; ****$P$<0.0001.

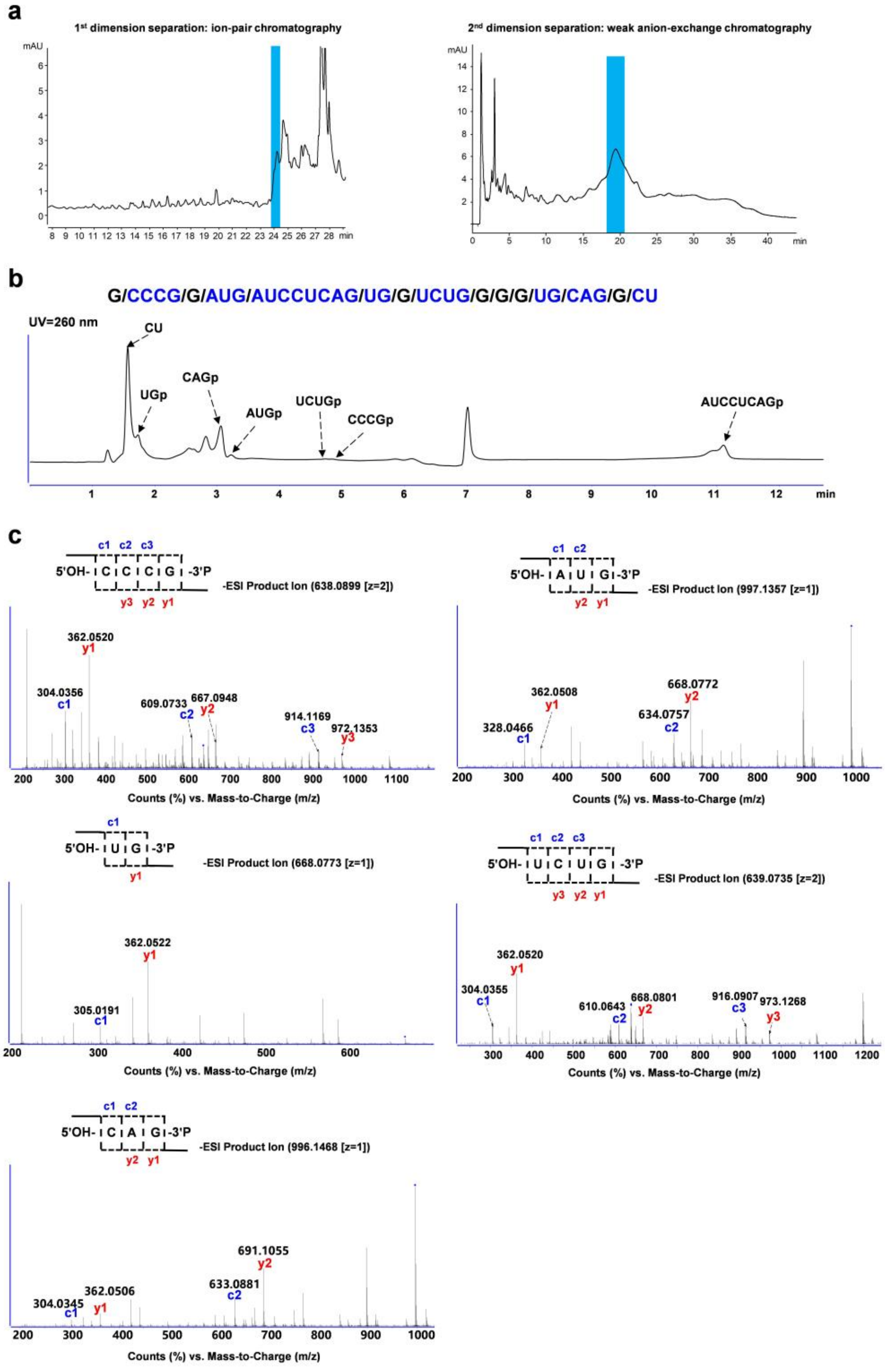


**Figure S2. Biological function of t-half fraction from aged mice and separation and characterization of 5′-tRNA$^{Sec(NCA)}$ half**. **a**, Two-dimensional liquid chromatographic chromatograms for purification of 5′-tRNA$^{Sec(NCA)}$ half. **b**, Annotations of RNase T1 digestions and sequence mapping of 5′-tRNA$^{Sec(NCA)}$ half. **c**, Detailed MS/MS spectra of RNase T1 digestion products of 5′-tRNA$^{Sec(NCA)}$ half.

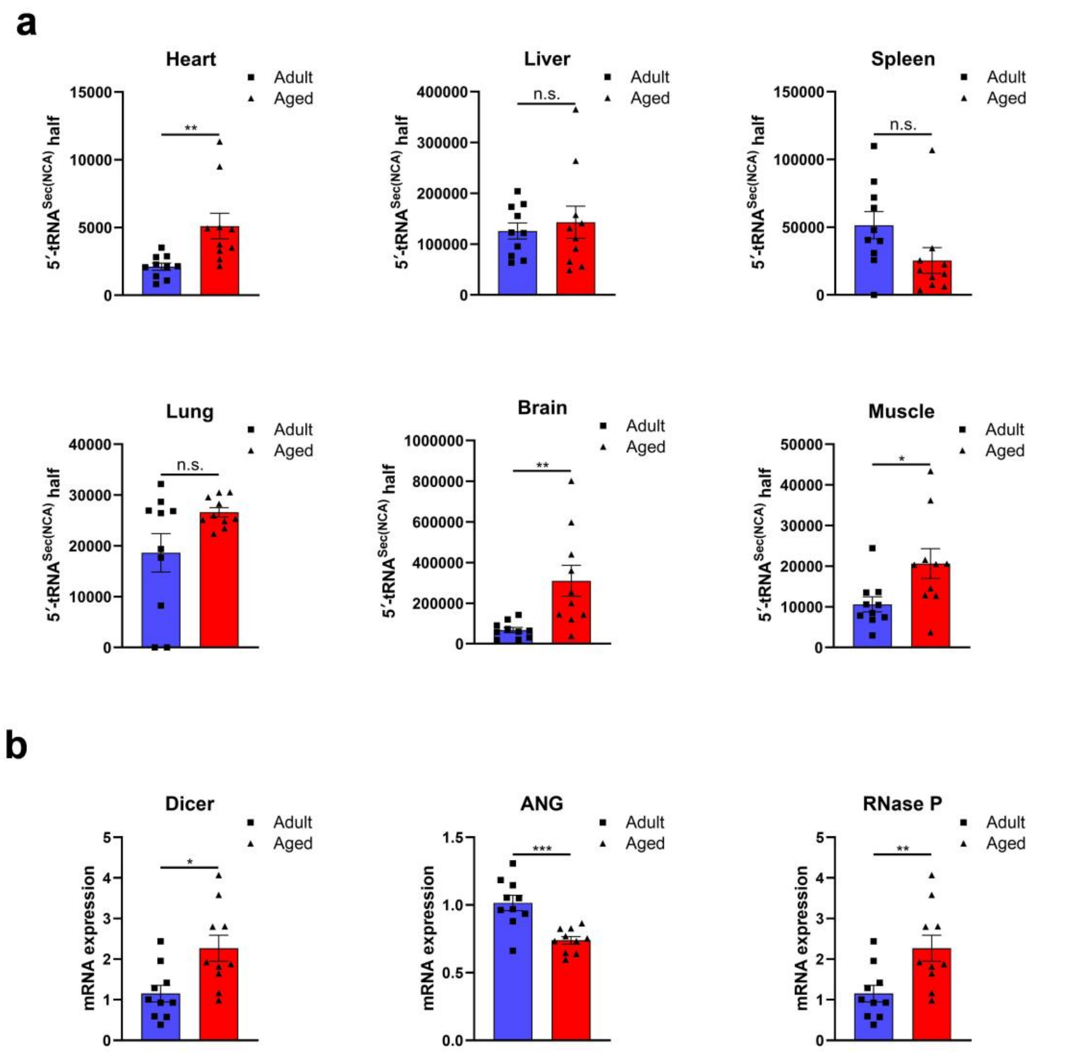


**Figure S3. Expressions of 5′-tRNA$^{Sec(NCA)}$ half and its potential cleavage-related ribonucleases during aging**. **a**, 5′-tRNA$^{Sec(NCA)}$ half expressions in major organs except for kidney from adult and aged mice determined by LC-MS analysis. **b**, mRNA expressions of potential ribonucleases for tRNA cleavage in kidneys from adult and aged mice. Data presented in all figures are presented as mean ± S.E.M. Statistical test: two-tailed unpaired t-test. **P*<0.05; ***P*<0.01; ****P*<0.001; *****P*<0.0001; n.s., non-significant.

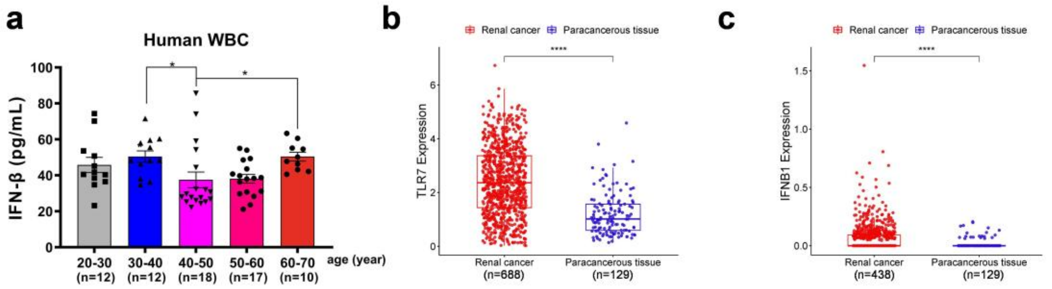


**Figure S4. Expressions of TLR7 and IFN-β in clinical samples**. **a**, IFN-β expressions in plasma from healthy humans of different age determined by ELISA kit. Expression levels of (**b**) TLR7 and (**c**) IFNB1 (encoding IFN-β) in renal cancer and paracancerious tissue using TCGA data. Data presented in **a**, **b** and **c** are presented as mean ± S.E.M. Statistical test: two-tailed unpaired t-test. $*P<0.05$; $****P<0.0001$.

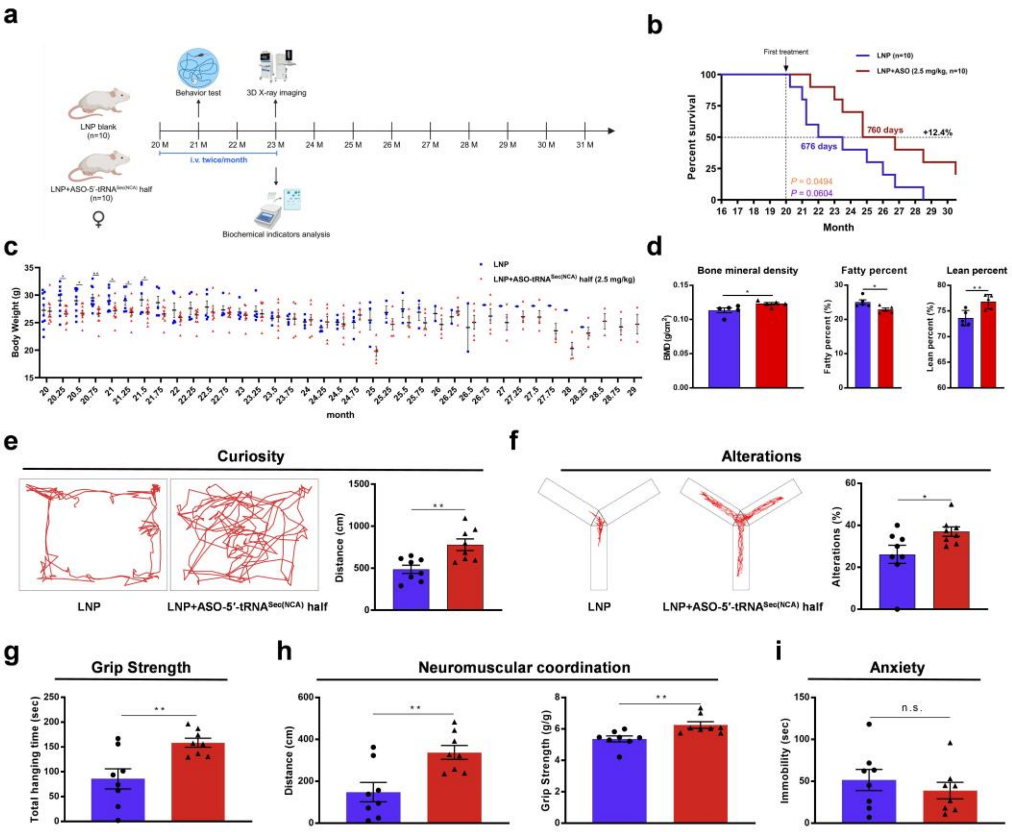

**Figure S5. ASO-5′-tRNA$^{Sec(NCA)}$ half extends the healthy lifespan of naturally aged female mice. a**, Experimental design of anti-aging test of ASO-5′-tRNA$^{Sec(NCA)}$ half from 20-month-old female Balb/c mice. Survival curves (**b**) and weight curve (**c**) of aged mice received LNP encapsulated ASO-5′-tRNA$^{Sec(NCA)}$ half treatment (n=10) or not (n=10). **d**, Bone mineral density, fatty percent and lean percent of mice. **e**, Movement of mice received ASO-5′-tRNA$^{Sec(NCA)}$ half treatment or not in the open-field test. **f**, Alteration rates of ASO-5′-tRNA$^{Sec(NCA)}$ half-treated mice or not in the Y maze test. Grip strength test (**g**) and rotarod test (**h**) of ASO-5′-tRNA$^{Sec(NCA)}$ half-treated mice or not. **i**, There was no significant difference in the immobility time between the two groups of mice. Data presented in **b** are presented as the log-rank test in orange and the Gehan-Breslow-Wilcoxon in purple. Data presented in **c**, **d**, **e**, **f**, **g**, **h**, and **i** are presented as mean ± S.E.M. Statistical test: two-tailed unpaired *t*-test. *$P$<0.05; **$P$<0.01; ***$P$<0.001; n.s., non-significant.

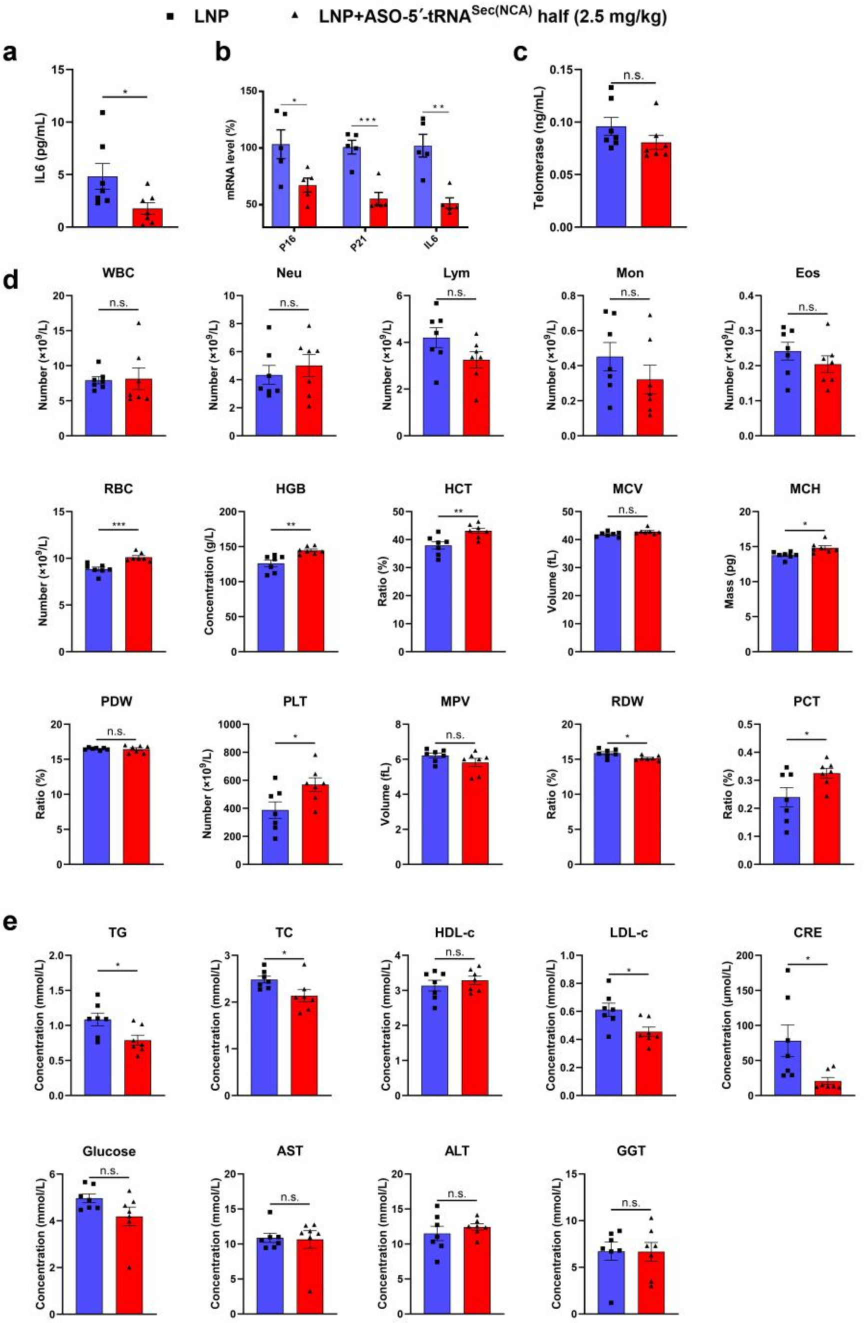


**Figure S6. ASO-5′-tRNA$^{Sec(NCA)}$ half ameliorates healthspan in naturally aged male mice**. **a**, ELISA detection of IL6 content in plasma of mice. **b**, qPCR analysis of P16, P21 and IL6 expressions in white blood cells isolated from mice. **c**, ELISA detection of telomerase content in plasma of mice. **d**, Complete blood analysis of WBC, Neu, Lym, Mon, Eos, RBC, HGB, HCT, MCV, MCH, PDW, PLT, MPV, RDW, AND PCT in aged mice. **e**, Evaluation of metabolism (TG, TC, HDL-c, LDL-c), glucose, kidney function (CRE) and liver function (AST, ALT, GGT) in aged mice. Data presented in all figures are presented as mean ± S.E.M. Statistical test: two-tailed unpaired t-test. **P*<0.05; ***P*<0.01; ****P*<0.001; n.s., non-significant.

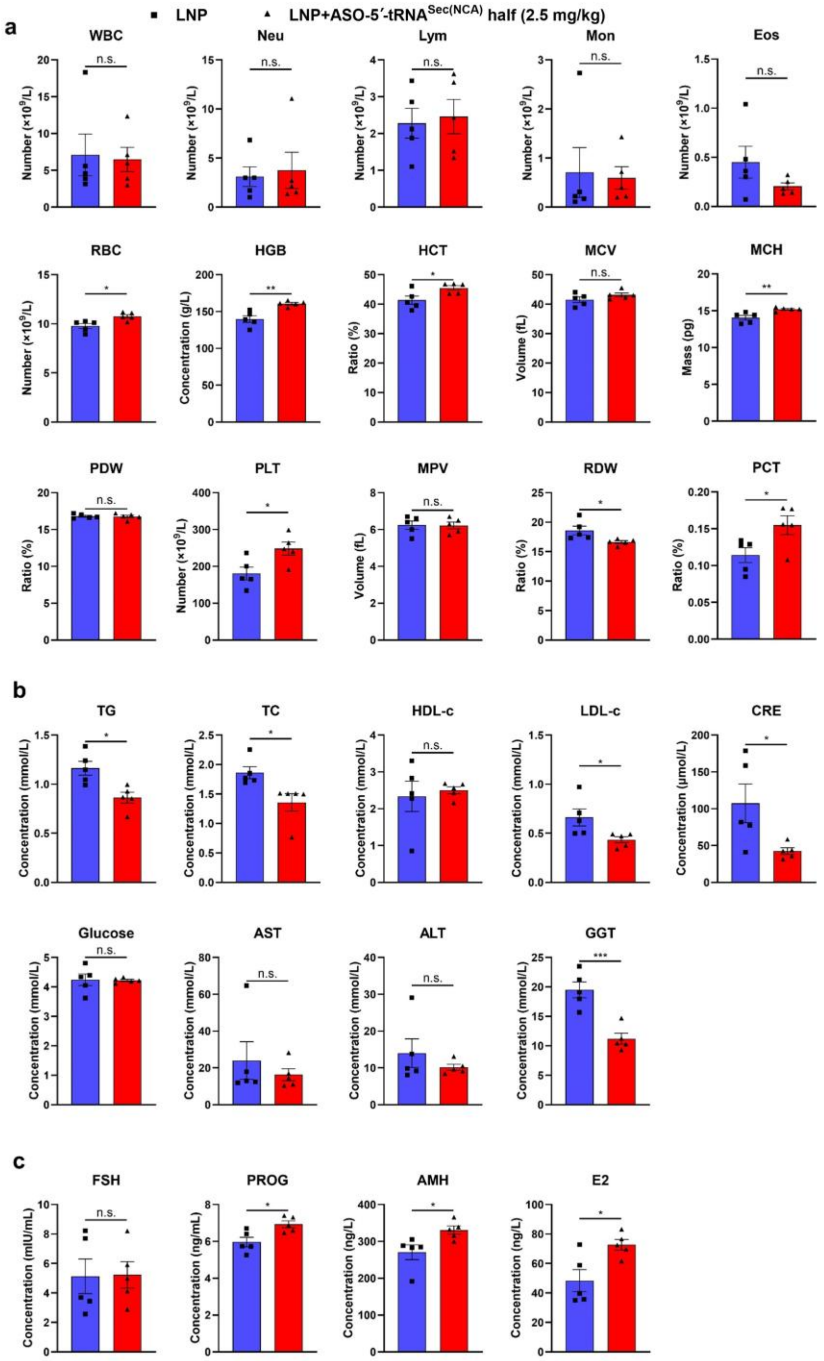


**Figure S7. ASO-5′-tRNA$^{Sec(NCA)}$ half ameliorates healthspan in naturally aged female mice**. **a**, Complete blood analysis of WBC, Neu, Lym, Mon, Eos, RBC, HGB, HCT, MCV, MCH, PDW, PLT, MPV, RDW, AND PCT in mice. **b**, Evaluation of metabolism (TG, TC, HDL-c, LDL-c), glucose, kidney function (CRE) and liver function (AST, ALT, GGT) from plasma. **c**, Evaluation of female reproductive hormones (FSH, PROG, AMH, E2) from plasma. Data presented in all figures are presented as mean ± S.E.M. Statistical test: two-tailed unpaired t-test. **P*<0.05; ***P*<0.01; ****P*<0.001; n.s., non-significant.

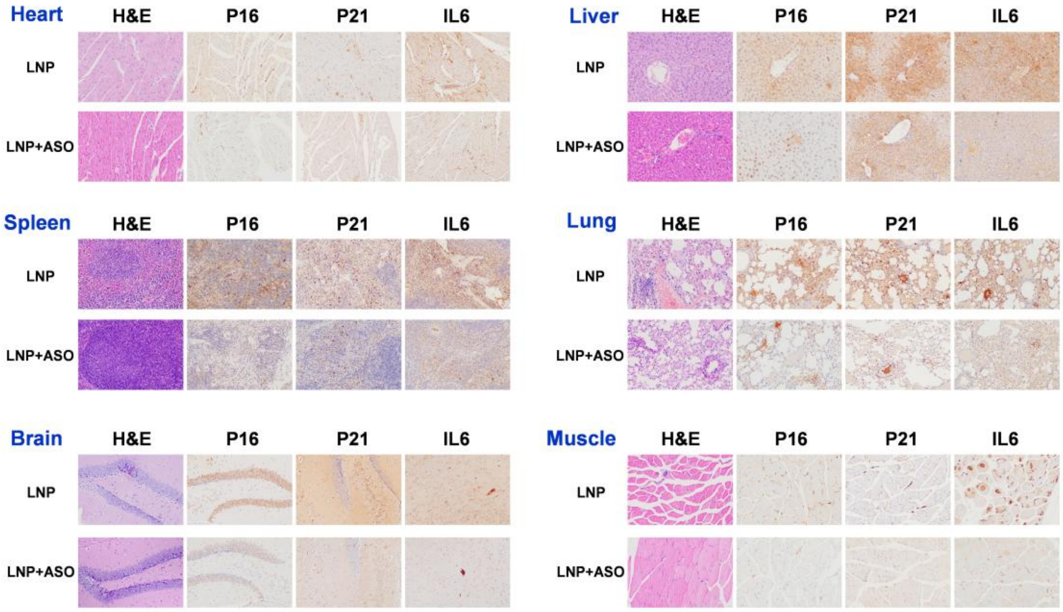


**Figure S8. H&E and IHC staining of P16, P21 and IL6 in major organs of naturally aged mice treated with ASO-5′-tRNA$^{Sec(NCA)}$ half**.
